# The INO80 Chromatin Remodeler Sustains Metabolic Stability by Promoting TOR Signaling and Regulating Histone Acetylation

**DOI:** 10.1101/184440

**Authors:** Sean L. Beckwith, Erin K. Schwartz, Pablo E. Garcia-Nieto, Devin A. King, Graeme J. Gowans, Ka Man Wong, Wei Yao, Tessa L. Eckley, Alexander P. Paraschuk, Egan Peltan, Laura R. Lee, Ashby J. Morrison

**Affiliations:** Department of Biology, Stanford University, Stanford, CA, USA; Current address: Department of Biological Sciences, Florida Atlantic University, Boca Raton, FL, USA; Current address: Department of Chemical and Systems Biology, Stanford University School of Medicine, Stanford, CA, USA

## Abstract

Chromatin remodeling complexes are essential for gene expression programs that coordinate cell function with metabolic status. However, how these remodelers are integrated in metabolic stability pathways is not well known. Here, we report an expansive genetic screen with chromatin remodelers and metabolic regulators in *Saccharomyces cerevisiae*. We found that, unlike the SWR1 remodeler, the INO80 chromatin remodeling complex is composed of multiple distinct functional subunit modules. We identified a strikingly divergent genetic signature for the Ies6 subunit module that links the INO80 complex to metabolic homeostasis, including mitochondrial maintenance. INO80 is also needed to communicate TORC1-mediated signaling to chromatin and maintains histone acetylation at TORC1-responsive genes. Furthermore, computational analysis reveals subunits of INO80 and mTORC1 have high co-occurrence of alterations in human cancers. Collectively, these results demonstrate that the INO80 complex is a central component of metabolic homeostasis that influences histone acetylation and may contribute to disease when disrupted.

## INTRODUCTION

Chromatin is a complex structure that is dynamically reorganized to facilitate DNA-templated processes such as transcription, chromosome segregation, DNA replication and DNA repair. Enzymes that restructure the chromatin environment are critical components of epigenetic maintenance and can contribute to disease when disrupted. Included among chromatin modifiers are enzymes that post-translationally modify histones and ATP-dependent chromatin remodelers that alter the position and composition of nucleosomes (Clapier & Cairns, 2009). Chromatin remodelers are evolutionarily conserved and regulate diverse processes required for normal cell function, organismal development and are mutated in a large fraction of cancers (Davis & Brachmann, 2003; de la Serna, Ohkawa, & Imbalzano, 2006).

Many remodelers are large multi-subunit complexes that can utilize the function of different subunits in a tissue-specific manner, allowing for cell-type specific regulation (Wu, 2012). In particular, different subunits of the evolutionarily conserved INO80 chromatin remodeling complex have demonstrated roles in diverse processes, such as transcription (Alcid & Tsukiyama, 2014; X Shen, Mizuguchi, Hamiche, & Wu, 2000; Xue et al., 2015), replication (Papamichos-Chronakis & Peterson, 2008; Shimada et al., 2008; Vincent, Kwong, & Tsukiyama, 2008), DNA damage responses (Attikum et al., 2004; Falbo et al., 2009; Morrison et al., 2004, 2007), telomere regulation (Yu et al., 2007), mitotic stability (Chambers et al., 2012; Ogiwara, Enomoto, & Seki, 2007), and metabolic homeostasis (Yao et al., 2016). These studies exemplify the functional diversity of the INO80 complex in different pathways (Morrison, 2017; Morrison & Shen, 2009; Poli, Gasser, & Papamichos-Chronakis, 2017), and suggest the partitioning of diverse functions among the subunits of the INO80 complex.

Individual subunits of the INO80 complex assemble within distinct structural modules along the ATPase subunit (Tosi et al., 2013; Watanabe et al., 2015). The Actin-related protein 8 (Arp8) module consists of Arp8, Arp4, Actin, Taf14 and Ies4. Arp4 and Arp8 are important for nucleosome recognition, ATP hydrolysis, and nucleosome sliding *in vitro* (Gerhold et al., 2012; Harata et al., 1999; Kapoor, Chen, Winkler, Luger, & Shen, 2013; Saravanan et al., 2012; Xuetong Shen, Ranallo, Choi, & Wu, 2003; Tosi et al., 2013). The N-terminal domain of the Ino80 ATPase assembles the Nhp10 module consisting of Nhp10, Ies1, Ies3, and Ies5, subunits that are less conserved among different species (Jin et al., 2005; Tosi et al., 2013). The Arp5 module is essential for chromatin remodeling activity and includes Arp5 and Ies6 subunits that are needed for ATP hydrolysis, nucleosome sliding, and histone exchange (Xuetong Shen et al., 2003; Tosi et al., 2013; Watanabe et al., 2015; Yao et al., 2015).

One recent example of specific subunit contribution to the function of the INO80 complex is the role of the Arp5 and Ies6 subunits in the regulation of metabolic gene expression (Yao et al., 2016). Specifically, Arp5 and Ies6 form an abundant subcomplex that can assemble into the INO80 complex, stimulating *in vitro* activity and activating carbon metabolism gene expression *in vivo.* Indeed, these results support an emerging model where chromatin modifying enzymes are responsive to the metabolic state of the cell and alter the chromatin landscape, thereby linking metabolic status to transcriptional responses (Gut & Verdin, 2013). Indeed, many chromatin-modifying enzymes use key metabolites as co-factors or substrates that can fluctuate in different metabolic conditions, including acetyl-CoA, nicotinamide adenine dinucleotide (NAD^+^), and ATP. For example, histone acetyltransferases (HATs) use nuclear acetyl-CoA in high glucose conditions to acetylate histones, creating a permissive state for transcription (Shi & Tu, 2015). Additionally, in low energy states, high NAD^+^ levels activate the SIRT1 histone deacetylase (HDAC) to deacetylate H3K9 at the rDNA loci, suppressing the highly energy-consuming process of ribosome biogenesis (Murayama et al., 2008). Lastly, chromatin remodeling enzymes use ATP to hydrolyze histone-DNA contacts as they reposition or restructure nucleosomes (Zhou, Johnson, Gamarra, & Narlikar, 2016).

In order to identify the *in vivo* mechanisms of INO80’s metabolic regulation, we created a genetic interaction map using the epistatic mini-array profile (EMAP) approach in *S. cerevisiae*. Genetic interactions can reveal how sets of proteins coordinate higher level biological functions and identify crosstalk between pathways and processes (Beltrao, Cagney, & Krogan, 2010). EMAPs have previously been used to decipher gene networks involved in the secretory system (Schuldiner et al., 2005), chromatin modification (Collins et al., 2007), and DNA damage responses (Bandyopadhyay et al., 2010; Guénolé et al., 2013).

We identified genetic interactions between many chromatin and metabolic regulators, in both nutrient rich media and metabolic stress conditions to reveal nutrient-specific interactions. Our work reveals that subunits of the INO80 complex are functionally diverse and define distinct genetic modules. Both the NHP10 and ARP5 genetic modules connect the INO80 complex to histone (de)acetylation. Interestingly, we find that the IES6 genetic module is relatively disconnected from the rest of the INO80 complex and genetically interacts with components of the Target of Rapamycin (TOR) pathway that are critical to the maintenance of metabolic homeostasis. These results place the INO80 complex as an important regulator of histone modification that is downstream of TOR signaling.

## RESULTS

### An EMAP of chromatin and metabolic regulators

Given the interplay between metabolism and epigenetics, we set out to comprehensively identify shared pathways in which chromatin and metabolic regulators function in *S. cerevisiae*. To do this, we conducted an EMAP of unstressed and metabolically challenged cells grown on rich media (untreated), rapamycin or ethanol, which generated approximately a quarter million interactions (Figure 1A **and Supplementary File 1**). Rapamycin inhibits the TORC1 complex, a master regulator of cellular growth (Loewith & Hall, 2011). Ethanol is a non-fermentable carbon source that requires cells to utilize oxidative phosphorylation, whereas yeast preferentially ferment glucose (Zaman, Lippman, Zhao, & Broach, 2008). We included a test library of 1536 alleles covering most major cellular processes, and significantly enriched for chromatin and metabolic regulators (Ryan et al., 2012). We used 54 query strains that cover several chromatin remodeling complexes, histone modifiers and metabolic signaling pathways (Figure 1B).

**Figure 1.**
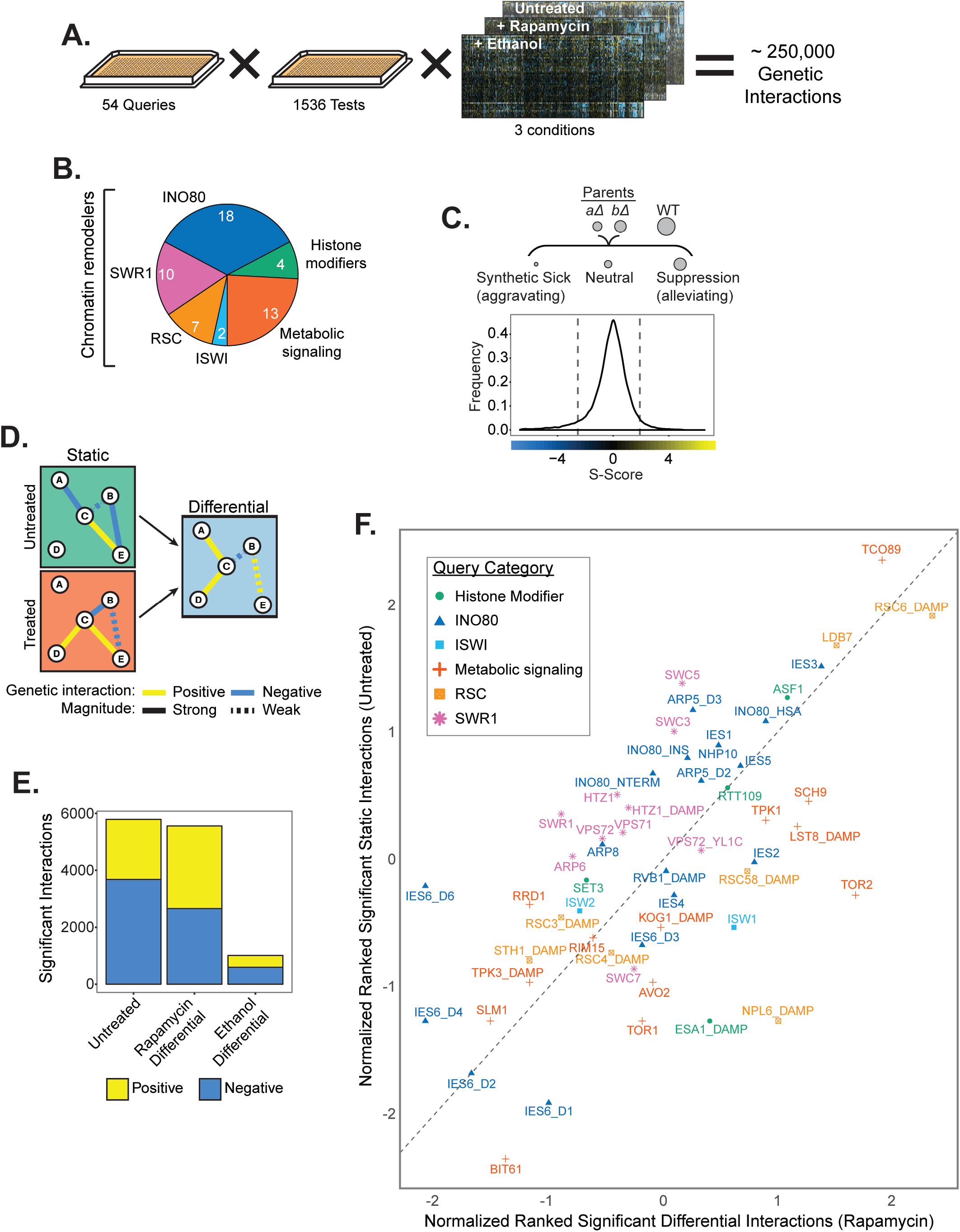
An epistasis map of chromatin and metabolic regulators. (**A**) Overview of EMAP including 54 query strains and a library of 1536 test strains, assayed in three growth conditions. (**B**) Composition of the query library by number of query strains; INO80, SWR1, RSC and ISWI are chromatin remodeling complexes. Histone modifiers include histone acetyl-transferases, histone deacetylases. Metabolic signaling genes include components of the TOR and PKA signaling networks. Numbers indicate the number of query strains in each category. (**C**) *Top*, Genetic interaction scores (S-score) are computed by comparing the observed fitness, inferred from colony size, of double mutants with the expected fitness, which is based on fitness of parental strains. A wild-type (WT) strain is shown for comparison. *Bottom*, the distribution of S-scores is shown for the untreated condition. Dashed lines indicate significance cutoffs of -2.5 and 2 for aggravating and alleviating interactions, respectively. (**D**) Hypothetical genetic interaction network indicating how the differential network is constructed by “subtracting” the untreated condition from treated condition. (**E**) The number of significant positive and negative interactions for each growth condition. (**F**) Plot of rankit normalized significant interactions by query gene in the untreated condition and the rapamycin differential condition. Color and shape indicate query gene category. Dashed line indicates y=x reference line. Significant interaction tallies are included in Supplementary File 2.

Our analyses also included deletions of all INO80’s unique subunits and domain mutants of *INO80, ARP5* and *IES6* (see *Materials and Methods*) because complete deletion resulted in inconsistent colony growth in the EMAP process (data not shown), thus confounding our ability to confidently determine genetic interactions. The resulting mutants disrupted the Arp8, Arp5 and Nhp10 structural modules of the INO80 complex (Figure 1 – **figure supplement 1**).

Genetic interactions (S-scores) were calculated from the fitness of double mutants (Figure 1C, Figure 1 – **figure supplement 2**). Positive (suppression/alleviating) S-scores often reveal epistatic genetic relationships and indicate that the fitness of the double mutant was better than expected. Negative (synthetic sick/aggravating) S-scores usually identify compensatory pathways and indicate worse fitness than expected (Collins, Schuldiner, Krogan, & Weissman, 2006). Differential interaction networks for rapamycin and ethanol were assessed by comparing interactions in treated and untreated growth conditions (Figure 1D, Figure 1 – **figure supplement 3 and 4**), as previously described (Bandyopadhyay et al., 2010).

Over 5000 significant interactions were identified in both the untreated and rapamycin differential networks (Figure 1E **and Supplementary File 2**). In the presence of rapamycin, several TOR pathway genes, such as the TORC1 effector kinase *SCH9 and* TORC1 subunit *LST8* have increased number of significant interactions, indicating that the differential network is broad and effective at identifying TOR dependent genetic interactions (Figure 1F). Several subunits of the INO80 chromatin remodeling complex (*IES2, IES4, IES6)* also have increased number of interactions in the rapamycin differential network, supporting a metabolic role for INO80. In contrast, the ethanol differential network yielded fewer genetic interactions and only a few query strains have increased significant interactions, suggesting a less dramatic reorganization of the genetic interaction landscape upon ethanol treatment than in response to rapamycin (Figure 1E, Figure 1 – **figure supplement 5**). Interestingly, four of the top five query strains with the most significant interactions in the ethanol differential condition were subunits of the INO80 complex (**Supplementary File 2**). As observed before, *arp5Δ* and *ies6Δ* mutants have higher growth rates than expected on ethanol, presumably because these mutants have increased respiratory capacity (Yao et al., 2016). These genetic results further highlight a critical function for INO80, and the Arp5-Ies6 module, as an interaction hub for cellular response to ethanol.

### Distinct genetic organization of the INO80 and SWR1 complexes

We first used our EMAP data to comprehensively map the functional modules within the INO80 complex by correlating the interaction profile of each query subunit across the test library in untreated growth conditions (Figure 2A). Using this method, we found that INO80 subunits were organized into 4 genetic modules, which were also independently identified in principal component analysis (PCA) when pairwise correlations were k-means clustered (Figure 2B). Notably, the Nhp10 structural module clustered genetically and included Nhp10, Ies1, Ies3, Ies5, and the Ino80 N-terminus on which the Nhp10 module assembles. Thus, the distinct *in vivo* function of the NHP10 genetic module is organized among the subunits that are physically associated. [Note, for clarity, genetic modules are denoted with all uppercase letters (e.g. NHP10 module) and structural modules are denoted with an uppercase first letter only (e.g. Nhp10 module)].

**Figure 2.**
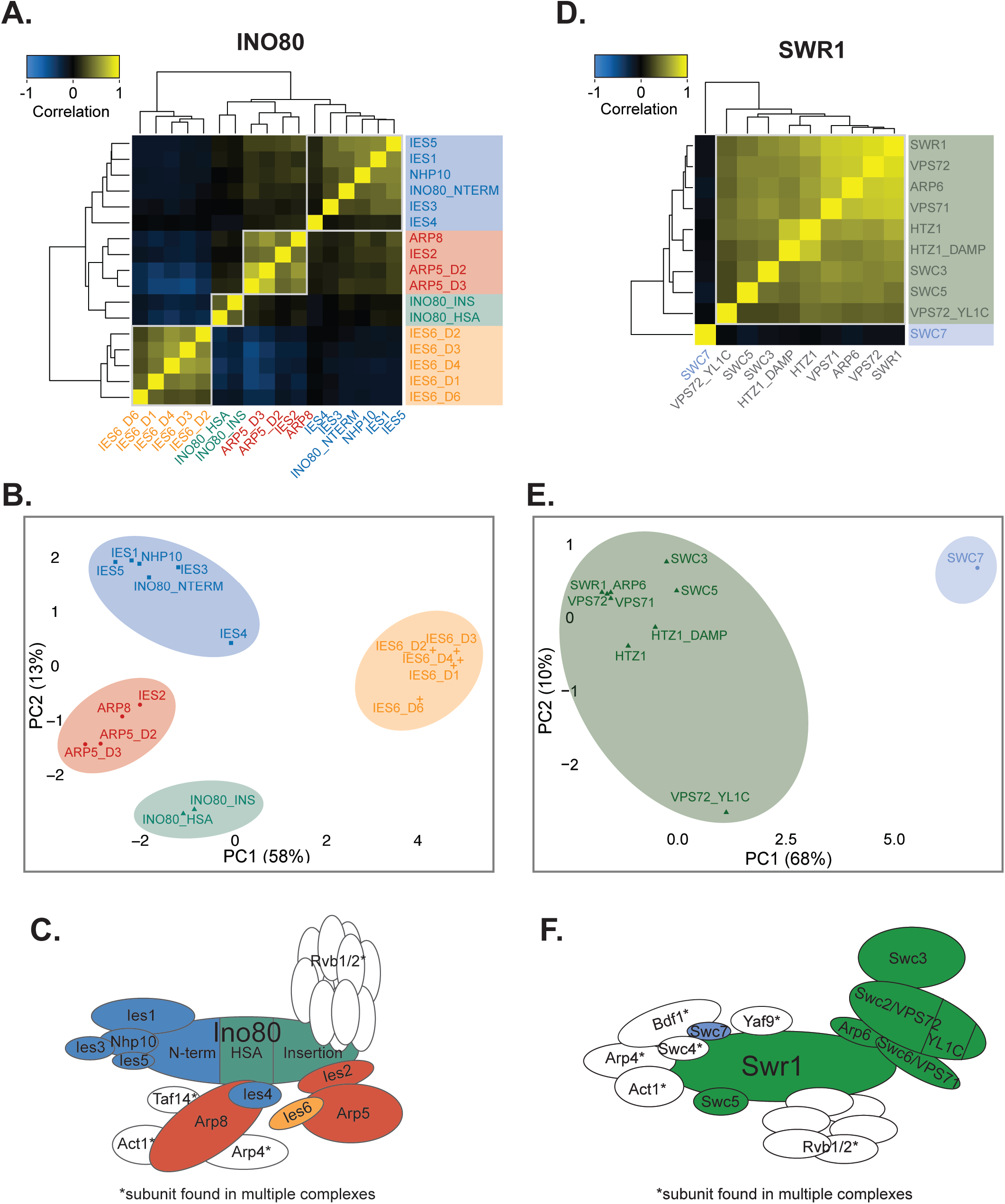
The INO80 complex is composed of distinct genetic modules. (**A**) Heatmap illustrating pairwise Pearson correlations between INO80 complex subunit query strains across the test library in the untreated static condition. Boxes outline genetic modules identified by hierarchical clustering and k-means analysis. Subunits that are not unique to the INO80 complex were omitted from the analysis. Mutants are knockout or domain deletions where indicated: *INO80* N-terminal (NTERM), insertion (INS), and HSA deletions; *ARP5* domain 2 and 3 (D2 and D3) deletions; and *IES6* domain 1, 2, 3, 4, and 6 (D1, D2, D3, D4, D6) deletions. (**B**) Principal component analysis (PCA) of Pearson correlations of INO80 complex subunit query strains as in (A). Colors indicate clustered genetic modules identified by k-means clustering (k=4). (**C**) Schematic illustrating the INO80 complex organized by known physical interactions (Tosi et al., 2013; Watanabe et al., 2015) with colors representing genetic modules of INO80 subunits identified in the untreated EMAP. (**D**) Heatmap of SWR1 complex subunit query strain Pearson correlations, as in (A). Mutants are knockout or domain deletions where indicated, *d*ecreased *a*bundance by *m*RNA *p*erturbation (DAmP) alleles are as described in Schuldiner et al., 2005. A Vps72 (Swc2) YL1-C domain mutant that is conserved in Ies6 was also included. (**E**) PCA of SWR1 strains as in (B), with k=2. (**F**) Schematic illustrating the SWR1 complex as in (C) based on structural studies (Nguyen et al., 2013).

However, other subunits of the INO80 complex assemble in genetic modules that are distinct from their structural modules. For example, Ies4 is structurally in the Arp8 module but was slightly more genetically similar to the NHP10 genetic module (Figure 2A and B). In addition, although Arp8 and Arp5 form separate structural modules, their genetic profiles are similar and constitute the ARP5 genetic module, which also includes *IES2*. Ies2 is needed for the Arp5 structural module to assemble with the INO80 complex (Yao et al., 2015), thus its *in vivo* function is tightly connected to Arp5 and is reflected in our genetic analysis.

The genetic signatures of the *INO80* helicase-SANT-associated (HSA) and insertion domain mutants were closely associated with each other and clustered closest to many subunits that assemble within those domains (Figure 2A and B). Namely, the HSA domain is required for association of Arp8 (Szerlong et al., 2008; Figure 1 – **figure supplement 1A and B**); and the insertion domain that splits *INO80*’s ATPase is required for the association of the Arp5 structural module (Yao et al., 2015). Most strikingly, all the domain mutants of *IES6* had genetic profiles that were dissimilar to the rest of the INO80 complex (Figure 2B). In fact, *IES6* mutant signatures anti-correlated with those in the ARP5 genetic module (Figure 2A). Given the physical association between Arp5 and Ies6 (Tosi et al., 2013; Yao et al., 2015, 2016), their divergent genetic profiles were extremely surprising. This genetic data suggests that, although Arp5 and Ies6 are physically coupled, they have some distinct and separable cellular functions. Figure 2C illustrates the INO80 complex genetic modules by color and are arranged according to previously identified structural modules (Tosi et al., 2013). INO80’s genetic architecture was not substantially changed in the rapamycin or ethanol EMAP (Figure 2 – **figure supplement 1A-D**).

In contrast to the INO80 complex, the SWR1 complex, another member of the INO80 chromatin remodeling subfamily (Mizuguchi et al., 2004), formed a strikingly cohesive genetic module (Figure 2D, E, and F). As before, analysis of non-unique subunits was not performed, such as several subunits that assemble in the N-terminal module of SWR1 (Nguyen et al., 2013) and are also found in the NuA4 acetyltransferase complex. Notably, our genetic analysis highlighted Swc7 as an outlier, the genetic profile of which did not correlate with other SWR1 subunits and formed a distinct module in PCA analysis and k-means clustering (Figure 2D and E). Figure 2F summarizes the genetic modules for SWR1, which are arranged according to previously identified structural modules (Nguyen et al., 2013). These genetic modules were largely preserved in the rapamycin and ethanol EMAP (Figure 2 – **figure supplement 1E-H**).

We next broadened our analysis to compare the SWR1 and INO80 complexes together to identify subunits that may facilitate cooperative or distinct function. Interestingly, the *SWC7* genetic profile was most similar to that of the *IES6* domain mutants (Figure 2 – **figure supplement 2**), suggesting that these subunits have common function that is distinct from both the SWR1 and INO80 complexes. In addition, the genetic profile of the *INO80* HSA and insertion domain mutants correlated with other SWR1 subunits and clustered with SWR1 subunits in PCA analysis. This suggests that the Ino80 ATPase and the SWR1 complex are involved in similar activities *in vivo.* Indeed, INO80 and SWR1 have many overlapping reported functions, including transcriptional regulation and genome maintenance (Gerhold & Gasser, 2014; Morrison & Shen, 2009). Additionally, high-resolution positional data shows similar binding profiles at +1 nucleosomes for both INO80 and SWR1 complex subunits (Yen, Vinayachandran, & Pugh, 2013), thus they may cooperatively regulate many genic loci.

Collectively, the EMAP results of the INO80 and SWR1 complex show very different genetic organization despite being of the same chromatin remodeling subfamily. Specifically, unique SWR1 C-terminus subunits are focused within similar *in vivo* functions, while the activities of the INO80 subunits are relatively more diverse and organized in distinct subunit modules. In addition, these analyses reveal that both Ies6 and Swc7 may not cooperatively function with their respective complexes, which may reflect independent activities for these subunits and/or regulatory roles that are not tested in the experimental conditions of this EMAP.

### Metabolic functions of the INO80 complex

In order to identify the cellular pathways in which the INO80 complex functions, we examined the function of genetically interacting test genes. Test genes with significant interactions to each genetic module were identified using DAVID functional annotation clustering analysis (Huang, Sherman, & Lempicki, 2009a, 2009b) (Figure 3A **and Supplementary File 3**), and individual biological process gene ontology enrichments are shown (Figure 3B **and Supplementary File 4**). Known functions of INO80 were captured in the EMAP, for example, chromatin modification, transcriptional regulation, and chromatin assembly are significantly enriched. Histone (de)acetylases and histone methylases were also identified as significant interactors, possibly due to cooperative functions as transcriptional regulators or direct effects by histone modifications on INO80’s activity. Mitotic functions, such as microtubule nucleation and mitotic spindle orientation were also identified in the INO80 genetic module, likely reflecting INO80’s role in chromosome segregation (Chambers et al., 2012; Hur et al., 2010).

**Figure 3.**
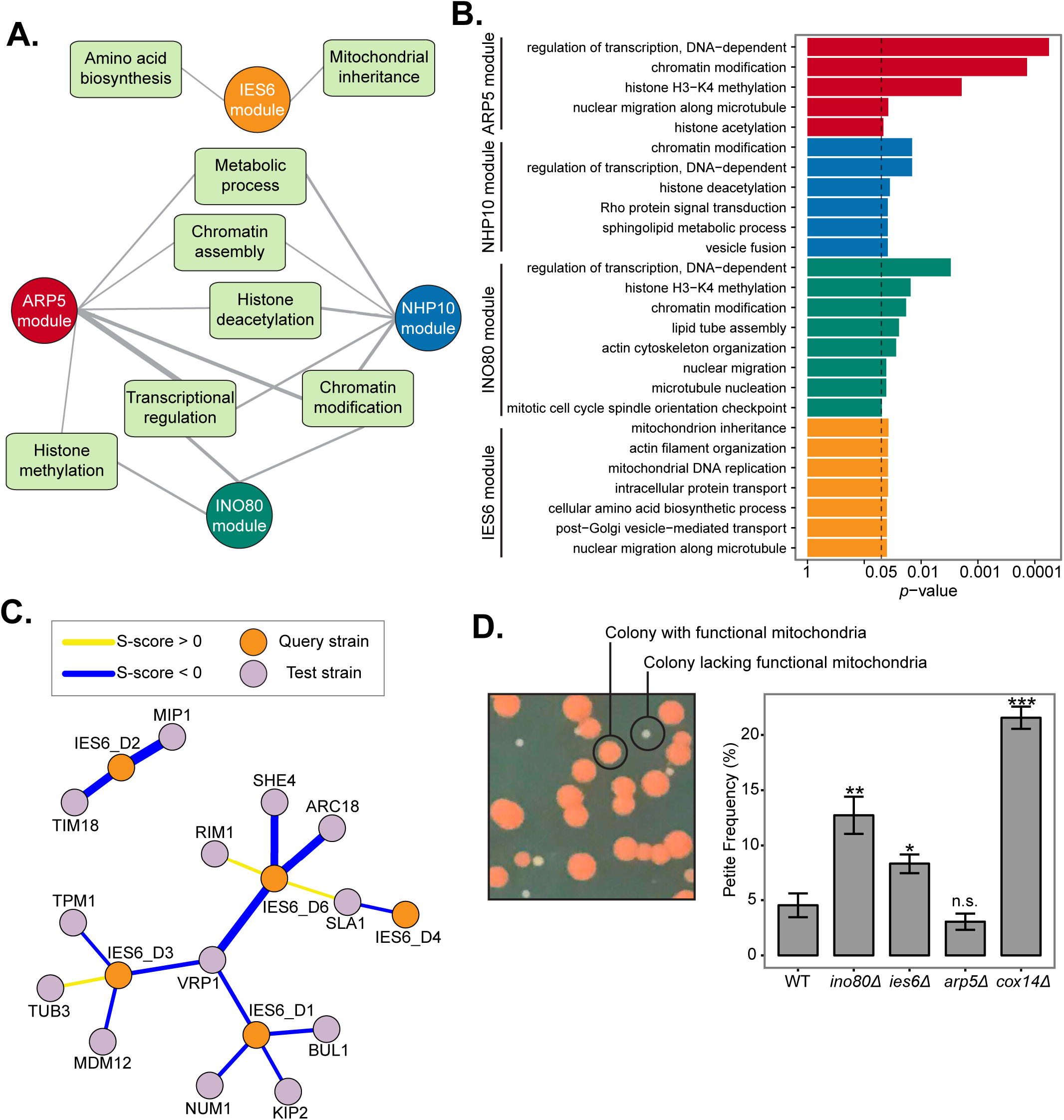
The IES6 genetic module is involved in mitochondrial maintenance. (**A**) Network diagram illustrating DAVID functional annotation clusters of significantly interacting test genes with each INO80 subunit query gene module identified in Figure 2. Line width indicates enrichment score, with a cutoff of ≥1.3 (-log_10_ p-value). Genes within each annotation are listed in Supplementary File 3. (**B**) FDR adjusted *p*-values of gene ontology (GO) enrichments (hypergeometric test, *p* < .05) of significantly interacting test genes with each INO80 subunit query gene module. The complete list of significant GO terms is found in Supplementary File 4. (**C**) Genetic interaction network between the IES6 genetic module and significantly interacting test genes found in the DAVID mitochondrial inheritance cluster. Line width indicates strength of S-score. (**D**) *Left*, representative image of yeast colonies overlaid with tetrazolium. Colonies founded by respiratory competent cells are large and red, “petite” colonies founded from respiratory deficient cells are smaller and white. *Right*, quantification of petite frequency in the indicated strains; deletion of *COX14* is known to increase petite frequency (Hess et al., 2009). Error bars represent standard error of the mean. Significance was determined using a Wilcoxon rank sum test from at least 8 independent measurements compared to wild-type.

Notably, the IES6 module did not overlap with the functional annotation clusters of the other modules and were significantly enriched in metabolic annotations, such as amino acid biosynthesis (Figure 3A), supporting previous findings of Ies6 in metabolic homeostasis (Yao et al., 2016). The only other significantly enriched annotation observed for the IES6 genetic module was mitochondrial inheritance. Corresponding test genes that interact with *IES6* domain mutants include several involved in cytoskeleton organization, such as *VRP1, ARC18* and *SLA1*, and mitochondrial membrane function and DNA replication, including *TIM18* and *MIP1* (Figure 3C).

To determine if Ies6 is directly involved in mitochondrial inheritance we utilized the previously established petite assay that examines the frequency of mitochondrial dysfunction (Hess et al., 2009). Deletion of the electron transport chain gene *COX14* served as a positive control and exhibited high petite frequency, as previously observed (Hess et al., 2009) (Figure 3D). Genetic deletions of *INO80* and of *IES6* exhibited high petite frequencies, while deletion of *ARP5* did not. As previously mentioned, the difference between the *ies6Δ* and *arp5Δ* mutants is surprising given that they physically interact each other (Yao et al., 2016). This assay further supports the notion that Ies6 and Arp5 have separable *in vivo* functions and demonstrate that the Ies6 subunit is needed for specific metabolic functions of the INO80 complex, including mitochondrial maintenance.

### INO80 is a regulator of histone acetylation

To further explore how INO80 functions among the other chromatin regulators in the EMAP, we examined the genetic interaction correlations between each query strain and the entire test library (Figure 4A). Interestingly, INO80 subunits were positively correlated with Rtt109 and Asf1, components of the H3K56 acetylase pathway that are important for genome stability (Collins et al., 2007) (Figure 4A, **blue panel**). Notably, H3K56ac has been reported to impact the histone variant exchange of Htz1 by INO80 and SWR1 *in vitro*, and high levels of H3K56ac leads to a decreased level of promoter-proximal Htz1 *in vivo* (Watanabe, Radman-Livaja, Rando, & Peterson, 2013). In order to investigate if these genetic similarities stem from shared transcriptional functions, we examined published microarray gene expression profiles (Lenstra et al., 2011) and found substantial correlations between INO80 subunits, Rtt109, and Asf1 (Figure 4B).

**Figure 4.**
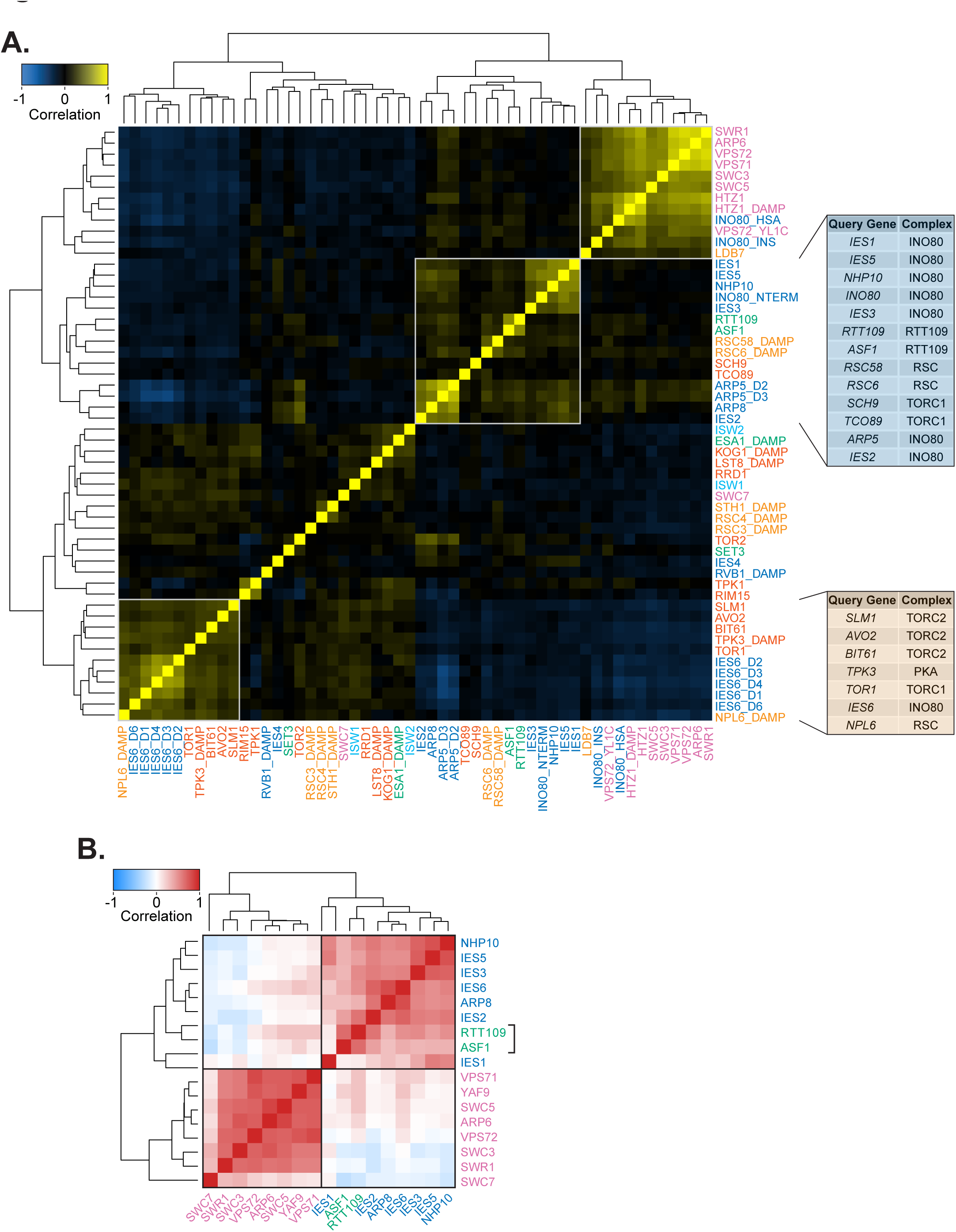
Genetic profiles of Rtt109 and metabolic regulators correlate with INO80. (**A**) Heatmap of Pearson correlation of all query strains in the untreated static condition. Label colors correspond to the query category annotated in Figure 1B. INO80 and SWR1 subunit mutants are described in Figure 2A, D. Boxes outline clusters identified by hierarchical clustering. *Right*, tables show the complex each query gene is found in for the INO80 and IES6 expanded genetic modules. (**B**) Heatmap of Pearson correlations of gene expression profiles from published microarray data (Lenstra et al., 2011) between deletion of subunits of the INO80 complex, SWR1 complex, *RTT109* and *ASF1*. All correlations between *RTT109, ASF1*, and INO80 subunits are significant, *p* < 0.001. Boxes indicate hierarchical clusters.

To explore whether Rtt109/Asf1 is a unique genetic interaction with INO80 or if INO80 is more broadly involved in histone modification status we next examined the genome-wide cooccupancy of the INO80 complex and all uniformly processed histone modification ChIP-seq datasets (see *Materials and Methods*, **Supplementary File 5**). We observed that Arp5 has the highest correlation with histone acetyl marks and anti-correlates with most histone methyl marks (Figure 5A). Corroborating the genetic interaction correlations between Rtt109, Asf1, and INO80 subunits, H3K56ac significantly correlates with Arp5 genome-wide (r = 0.53).

**Figure 5.**
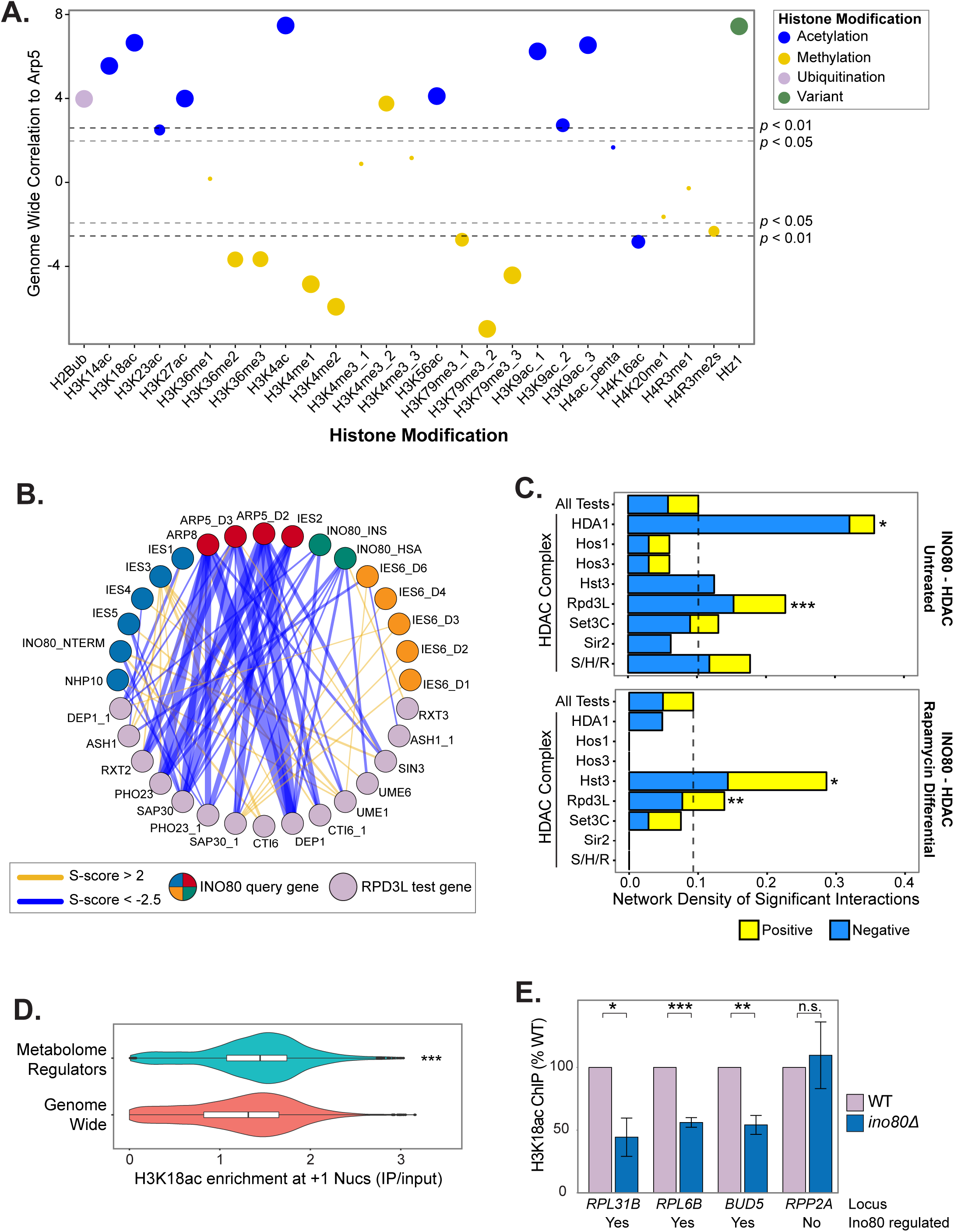
INO80 is a regulator of histone acetylation. (**A**) Genome-wide correlation of occupancy between Arp5 and histone modifications, listed on X-axis, using uniformly processed ChIP-seq data (see *Materials and Methods*). Colors illustrate modification type and size corresponds to binned *p*-value. (**B**) Genetic interaction network between INO80 subunit query strains and significantly interacting Rpd3L subunit test strains in the untreated static condition. Line width indicates strength of S-score, INO80 queries are colored according to modules identified in Figure 2. (**C**) Bar chart of network density by positive or negative significant interactions of test strains in the histone deacetylates complexes (HDACs) in yeast and INO80 subunit query strains in untreated or rapamycin differential conditions. Dashed line indicates the network density of all test strains (All Tests) and serves as a background benchmark. S/H/R is Sum1/Hst1/Rfm1. Significance was determined by Monte Carlo randomization test. (**D**) Violin and box plots of +1 nucleosome H3K18ac levels show significant regulators of the metabolome (Mülleder et al., 2016) (adjusted *p*-value < 0.01) have high H3K18ac levels compared to genome wide (*p*-value < 4.4e-16 by Wilcoxon rank sum test; *p* = 1.0e-5 by Monte Carlo randomization test). (**E**) ChIP-qPCR of H3K18ac in wild-type (WT) and *ino80Δ* deletion strains at loci chosen by H3K18ac levels from published data (Weiner et al., 2015) and regulation of expression by Ino80 (Yao et al., 2016). Significance was determined by Students *t*-test from at least 3 biological replicates, error bars represent standard error of the mean. Below each loci label is noted whether the gene’s expression is Ino80 regulated.

We then investigated the genetic interactions between INO80 query subunits and histone acetyltransferase and deacetylase test genes to further understand the relationship between INO80 and histone (de)acetylation. We found that INO80 has the highest density of significant interactions with the Rpd3L and HDA1 histone deacetylases in untreated, nutrient rich, conditions (Figure 5B and C). Rapamycin treatment did not significantly alter the genetic interactions between INO80 and HDA1 (Figure 5C, **bottom panel**). However, the interaction network density with Rpd3L was significantly enriched in the differential EMAP. This result is consistent with previous findings that Rpd3L, not HDA1, regulates histone deacetylation at TORC1-responsive genes (Humphrey, Shamji, Bernstein, & Schreiber, 2004; Rohde & Cardenas, 2003). Additionally, the network density between INO80 and both the Hst3 sirtuin histone deacetylase and SAGA histone acetyltransferase significantly increases in the presence of rapamycin (Figure 5C, **bottom panel** for Hst3 and data not shown, *p* = 0.00275, for SAGA). Both SAGA and Hst3 regulate the acetylation status of shared histone targets, the deacetylation of which is suppressed by TORC1 (Workman, Chen, & Laribee, 2016). Thus, the INO80 complex likely functions with different (de)acetylases depending on the metabolic environment.

To further investigate INO80’s maintenance of histone acetylation, we directly tested the effect of Ino80 loss on H3K18 acetylation (H3K18ac). We chose H3K18 because it is TORC1-responsive and deacetylated by Rpd3L and Hst3 (Workman et al., 2016). Additionally, H3K18ac and Arp5 have similar average distributions around +1 nucleosomes genome-wide (Figure 5 – **figure supplement 1A**), thus are able to regulate the same genes. We also found high H3K18ac levels at the +1 nucleosome of genes that significantly regulate the yeast metabolome (Mülleder et al., 2016) (Figure 5D). Accordingly, genes with high H3K18ac at the +1 nucleosome are also highly enriched for metabolome regulators (Figure 5 – **figure supplement 1B**). H3K18ac likely serves as a proxy for several histone acetylations at metabolic loci, as H3K18 occupancy significantly correlates (median r = 0.90) with several other acetyl marks at the +1 nucleosome genome-wide (Figure 5 – **figure supplement 2A and B**). We found that following deletion of *INO80*, H3K18ac was significantly reduced at several INO80-regulated genes (Figure 5E). Collectively, these results indicate that the INO80 complex cooperates with histone (de)acetylases to enact TORC1-mediated gene expression responses.

### INO80 is an effector of TOR signaling

We found strong evidence to support the role of INO80 as a TOR effector, as subunits of both TOR complex 1 and 2 (TORC1 and TORC2, respectively) and Sch9 downstream signaling kinase have positively correlated genetic interaction profiles with INO80 subunits (Figure 4A, **tan panel**). Strikingly, 5 of the 6 genes that correlate with the IES6 genetic module are subunits of the TORC1/2 and PKA signaling pathways and form an expanded IES6 metabolic module. This IES6 metabolic module was significantly enriched in test genes involved in many metabolic processes, such as amino acid biosynthesis, mitochondrial signaling, and intracellular transport (Figure 5 – **figure supplement 3A and Supplementary File 6**). Treatment with rapamycin markedly reduced the strength of the genetic interaction correlations for the expanded IES6 metabolic module, confirming that the genetic interactions between query and test genes are specific to nutrient-rich conditions and significantly reduced when TORC1-signaling is inhibited (Figure 5 – **figure supplement 3B**). INO80 and TORC1 have a highly connected genetic interaction network, both in rich media (Figure 6) and even more significantly in the rapamycin differential condition (Figure 6 – **figure supplement 1**), further supporting the interplay between INO80 and the TORC1 pathway.

**Figure 6.**
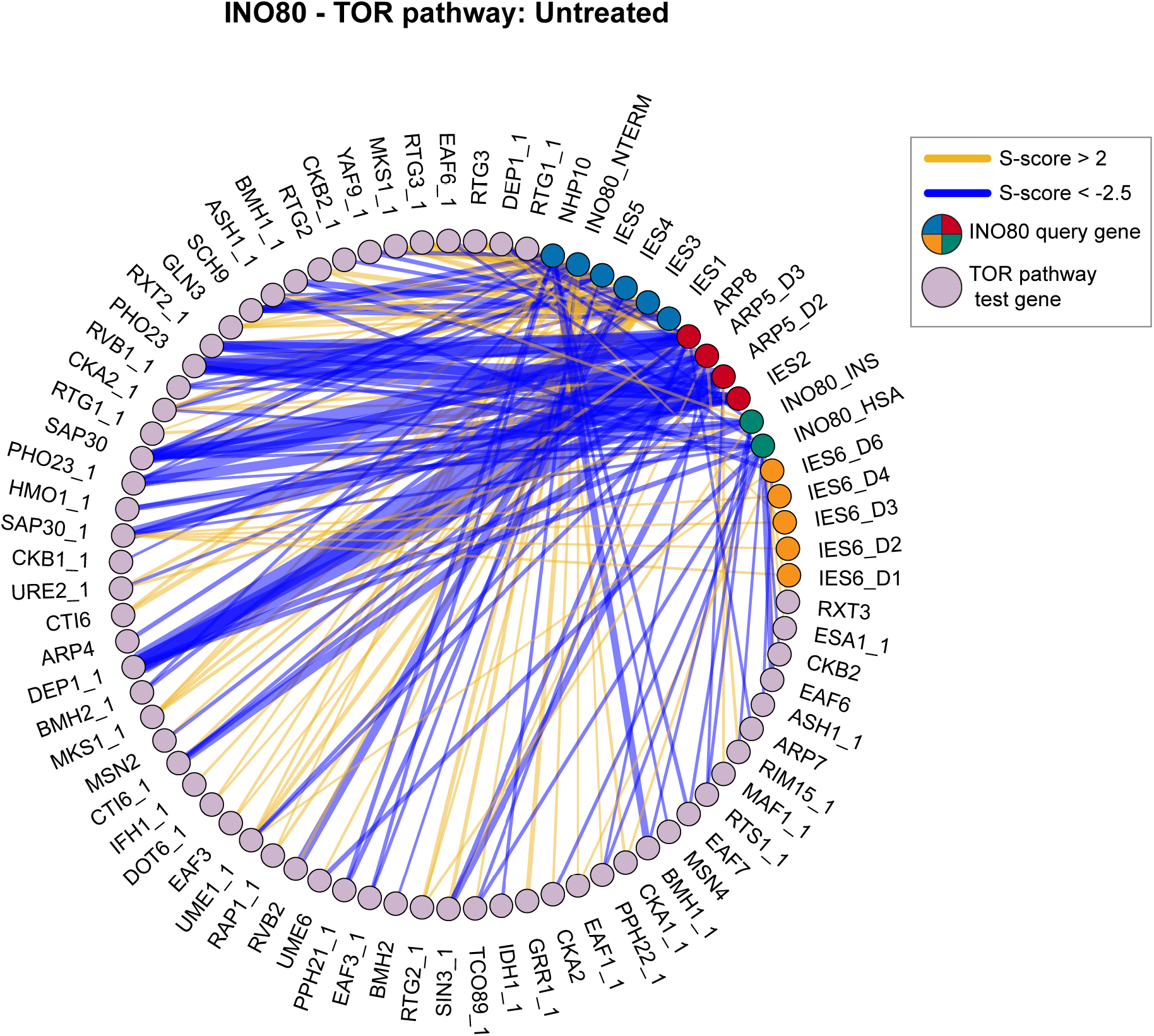
INO80 and the TOR pathway have a highly connected genetic interaction network. Genetic interaction network between INO80 subunit query strains and significantly interacting TOR pathway test strains in the untreated static condition. Line width indicates strength of S-score, INO80 queries are colored according to modules identified in Figure 2. Network density is significantly high, *p*-value = 1.1e-3 by Monte Carlo randomization test.

These results prompted us to further investigate how INO80 functions with TORC1 signaling. Interestingly, RNA-sequencing comparisons between rapamycin-treated cells and *ino80Δ* or *arp5Δ* mutant strains found similarities in gene expression profiles (r = 0.34, 0.31, respectively) (Figure 7A). A similar correlation (r = 0.34) was found comparing microarray expression data between *ies2Δ* (Lenstra et al., 2011) and rapamycin-treated cells (Urban et al., 2007). In fact, of the over 150 chromatin mutants analyzed (Lenstra et al., 2011), the expression profile of *ies2Δ* has the third highest correlation with rapamycin-treated cells (data not shown). Loss of *INO80* mimics many gene expression effects of rapamycin treatment, albeit to a lesser degree, including nitrogen metabolism, Msn2/4 stress response genes, and ribosome biogenesis (Figure 7B **and Supplementary File 7**). The expression of TORC1-responsive signaling and downstream transcription factors are similarly misregulated in both *ino80Δ* and rapamycin-treated cells.

**Figure 7.**
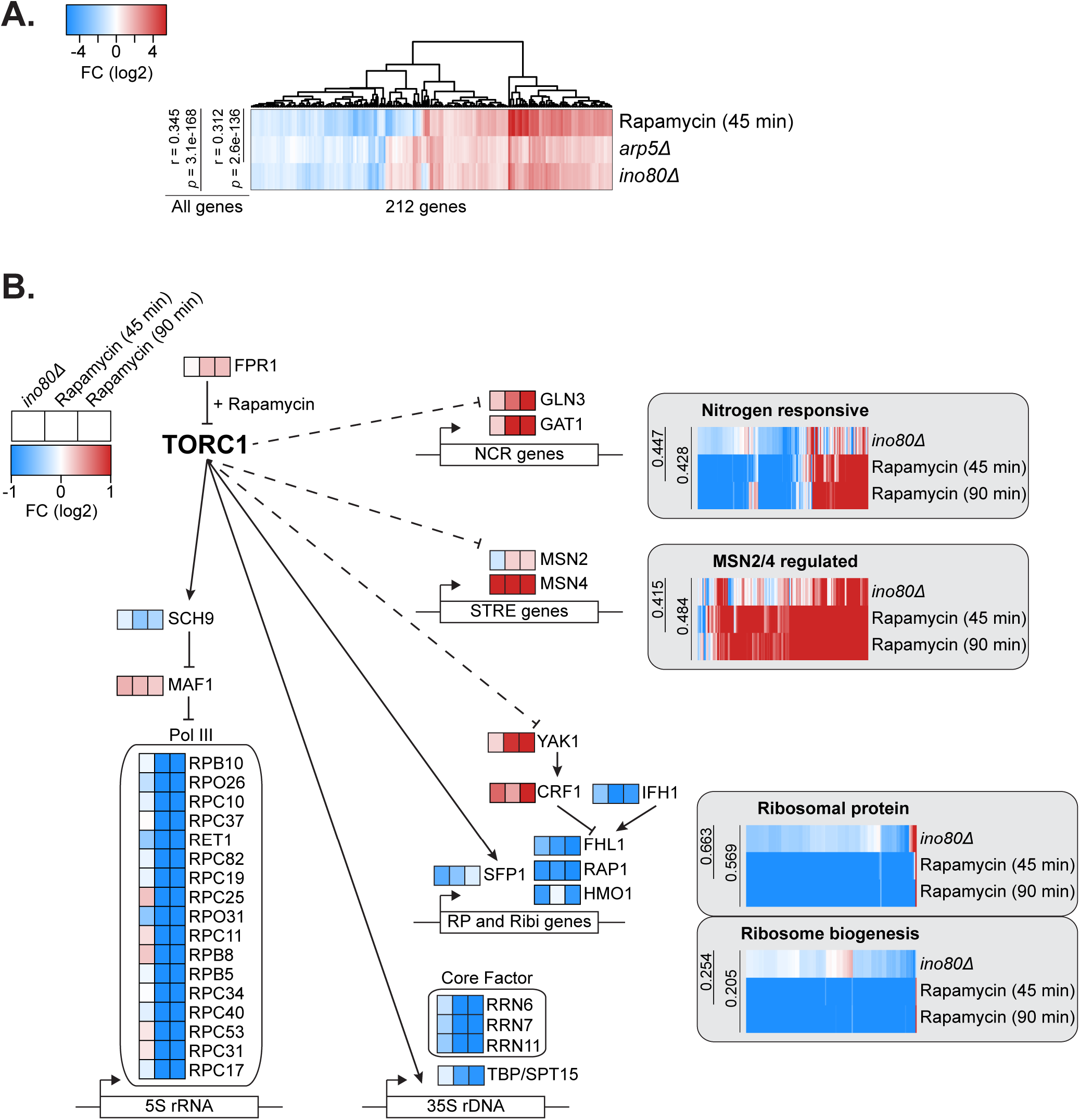
INO80 regulates the expression of key TOR signaling effectors. (**A**) Log-transformed Z-scores of expression fold-change (FC) between untreated and treated (30nM rapamycin for 45 minutes) wild-type cells or indicated knockout strains. Genes with at least a 1.5 fold-change are plotted. Pearson correlations and *p*-values are shown for all genes (>6000) regardless of fold-change difference. (**B**) Diagram of key genes involved in the TORC1 regulation of nitrogen source quality responsive genes (Scherens, Feller, Vierendeels, Messenguy, & Dubois, 2006), MSN2/4 regulated stress response genes (Gasch et al., 2000), ribosomal protein (RP) genes, and ribosome biogenesis genes (Jorgensen, 2004). Log-transformed expression fold-change is shown comparing untreated wild-type cells and rapamycin treated (45 and 90 minutes) or knockout strains as indicated. Gene lists are found in Supplementary File 7.

We also observed that *ino80Δ* cells were much less responsive to rapamycin treatment, which may result from compensatory mechanisms that emerge as a result of constitutively diminished TORC1-mediated transcription. Specifically, following rapamycin treatment, TORC1-responsive ribosomal protein (RP) gene expression in *ino80Δ* mutants is not decreased to the same degree as in wild-type cells (Figure 8A). Additionally, TORC1-dependent phosphorylation of Rps6, a ribosome component, persists in *ino80Δ* mutants following rapamycin treatment (Figure 8B). We also found that in growth assays, *ies6Δ* and *ino80Δ* mutants are resistant to rapamycin treatment (Figure 8C). Collectively, these observations demonstrate that loss of INO80 function results in persistent inability to transmit TORC1 signaling to chromatin and the creation of rapamycin refractory cells.

**Figure 8.**
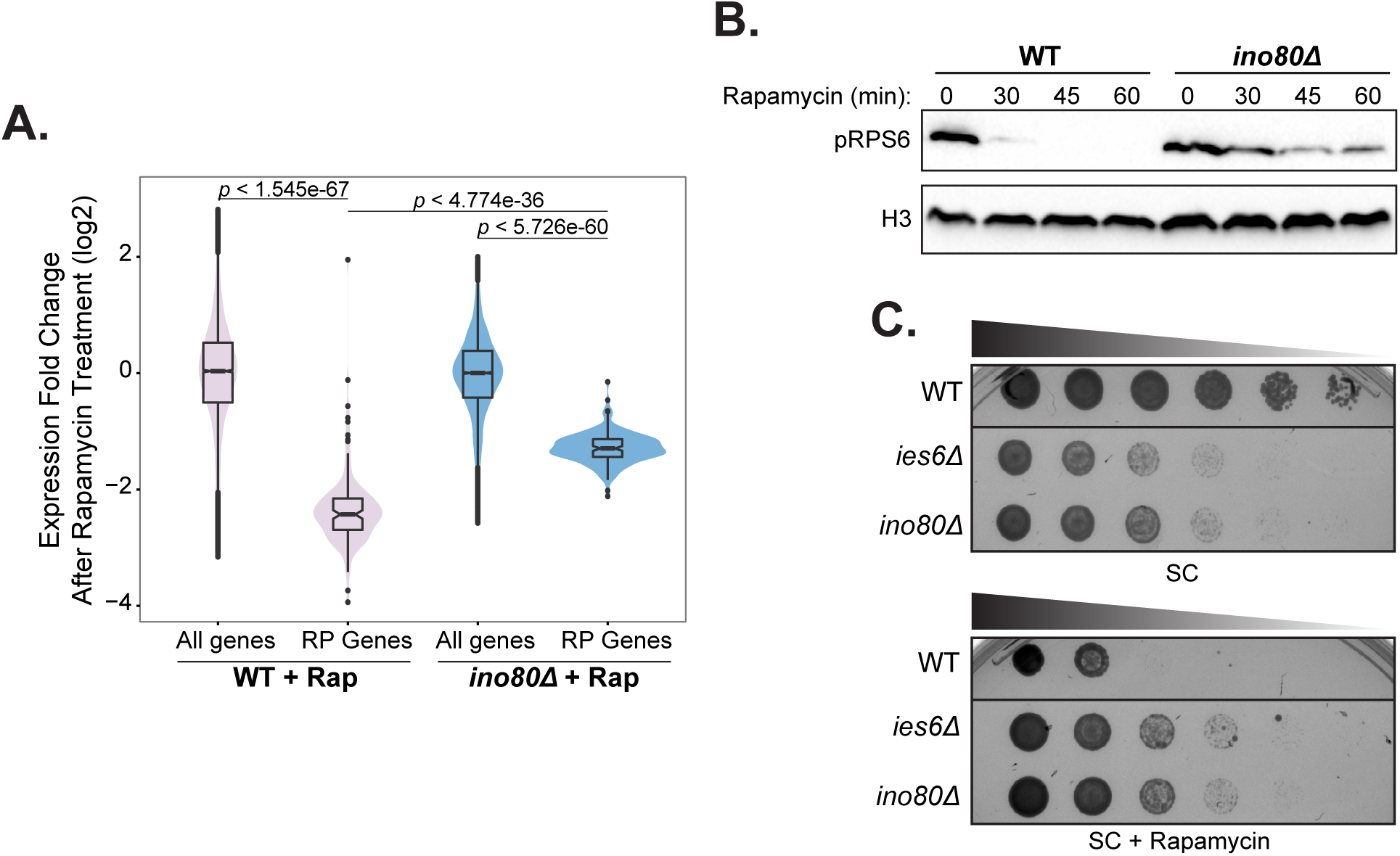
INO80 is an effector of the TORC1 pathway. (**A**) Violin and box plots of log-transformed expression fold-change after 45 minutes of 30 nM rapamycin (Rap) treatment compared to untreated cells in the indicated strains. The top and bottom 3% of genome wide responses were excluded for plotting. Significance was determined using a Wilcoxon rank sum test with the all genes. (**B**) Western analysis of phospho-Rps6 (pRps6) reduction following 30 nM rapamycin treatment for indicated minutes (min) in wild-type (WT) and *ino80Δ* strain. Histone H3 (H3) is a loading control. (**C**) Fitness assay of knockout strains compared to wild-type (WT). Serial dilution (1:5) of strains were grown at 30 °C on synthetic complete (SC) media with 0 or 5nM rapamycin.

## DISCUSSION

In this report, we examine an expansive genetic map to identify the functional composition of the INO80 complex. Unlike that of SWR1 unique subunits, the IN080 complex is genetically diverse and partitioned among several distinct modules. Partial function of the INO80 complex is constrained within structural modules (Tosi et al., 2013), such as the Nhp10 module, the subunits of which have cohesive genetic signatures. However, unexpected diversity is found with the Ies6 subunit, which forms a distinct genetic module that is anti-correlated with other INO80 subunits, including Arp5, its physical partner (Tosi et al., 2013; Watanabe et al., 2015; Yao et al., 2016). Interestingly, these unique Ies6 genetic interactions are enriched in metabolic functions and reveal previously unknown activities for the INO80 complex in mitochondrial maintenance and TOR signaling.

The role of Ino80 and Ies6 in mitochondrial inheritance (Figure 3D) may be indicative of a broader role for INO80 and/or Ies6 in the organization of organelles via the cytoskeleton. INO80 subunits genetically interact with genes involved in microtubule nucleation, actin cytoskeleton organization, and vesicle fusion (Figure 3B). Furthermore, in another genetic study, INO80 was connected to multivesicular body (MVB) sorting, cell polarity and morphogenesis, and cytokinesis (Costanzo et al., 2016).

Importantly, our genetic data has uncovered a strong connection between INO80 and TORC1, a rapamycin sensitive complex that is a master regulator of cell growth in yeast, plants and animals (Loewith & Hall, 2011). TORC1 signaling is active in nutrient rich conditions and promotes ribosome biogenesis while repressing cellular stress responses (Wei & Zheng, 2011; Figure 7B). INO80 and TORC1 have shared functions both in nutrient rich and rapamycin stress conditions, as indicated by correlated genetic profiles (Figure 4A; Figure 1 – **figure supplement 2 and 3**) and direct genetic interactions between INO80 and the TORC1 signaling pathway (Figure 6; Figure 6 – **figure supplement 1**). INO80 subunits are hub genes, that is highly connected nodes, in our rapamycin differential network, supporting a central role for INO80 in responding to rapamycin treatment. Additionally, similar transcriptional profiles are observed in *ino80*Δ mutants and cells treated with rapamycin (Figure 7A). Collectively, these data suggest that INO80 is needed to communicate TORC1-mediated growth signaling to chromatin.

One way in which INO80 can facilitate TORC1-dependent gene expression is by regulating histone acetylation status, thus transcriptional potential. Our study finds that INO80 genetically interacts with the acetyltransferases Rtt109 and SAGA, and with several rapamycin-responsive deacetylases, including Rpd3L and Hst3 (Figure 4A and B; Figure 5C). Interestingly, both Rpd3L and acetylated H3K56, the product of Rtt109 acetylation, are in the TORC1 signaling pathway (Chen, Fan, Pfeffer, & Laribee, 2012; Huber et al., 2011; Humphrey et al., 2004). The genome occupancy of Arp5 and acetylated H3K56 correlate, as do many histone acetyl marks, and loss of *INO80* reduces histone acetylation at metabolic loci (Figure 5A and E). Thus, INO80 may function to promote histone acetylation on growth genes downstream of TORC1.

Histone acetylation is also intimately linked to metabolic status, as it requires the metabolic intermediate acetyl-CoA. High levels of histone acetylation are present on genes that regulate the metabolome (Figure 5D; Figure 5 – **figure supplement 1B and 2A**), perhaps reflecting a feedback loop, whereby expression of metabolome regulators promotes acetyl-CoA production, which subsequently increases histone acetylation and gene expression. Thus, changes in metabolite availability could signal environmental conditions that are translated through chromatin. Future research will be needed to determine the role of INO80 and other chromatin remodelers that link metabolic status to epigenetic programming.

However, it is known that the consequences of deregulated metabolic signaling often result in disease. Indeed, energy metabolism alterations are a major contributing factor for many pathologies, including cancer, cardiovascular disease, and diabetes, which together account for two-thirds of all deaths in industrialized nations. For example, the mTOR signaling pathway is often constitutively active in cancer, promoting growth signaling irrespective of metabolic environments (Laplante & Sabatini, 2009). In this study, we find that the INO80 complex is needed to enact TORC1-responsive transcriptional programs. As both TORC1 and INO80 are conserved from yeast to humans, we investigated overlapping mutational signatures in cancer patient datasets. Indeed, we observed a high co-occurrence of alterations in subunits of hINO80 and mTORC1 in a wide range of human cancers (Figure 9 **and Supplementary File 8**), suggesting that abrogation of both INO80 and mTORC1 may lead to the metabolic dysregulation that contributes to carcinogenesis.

**Figure 9.**
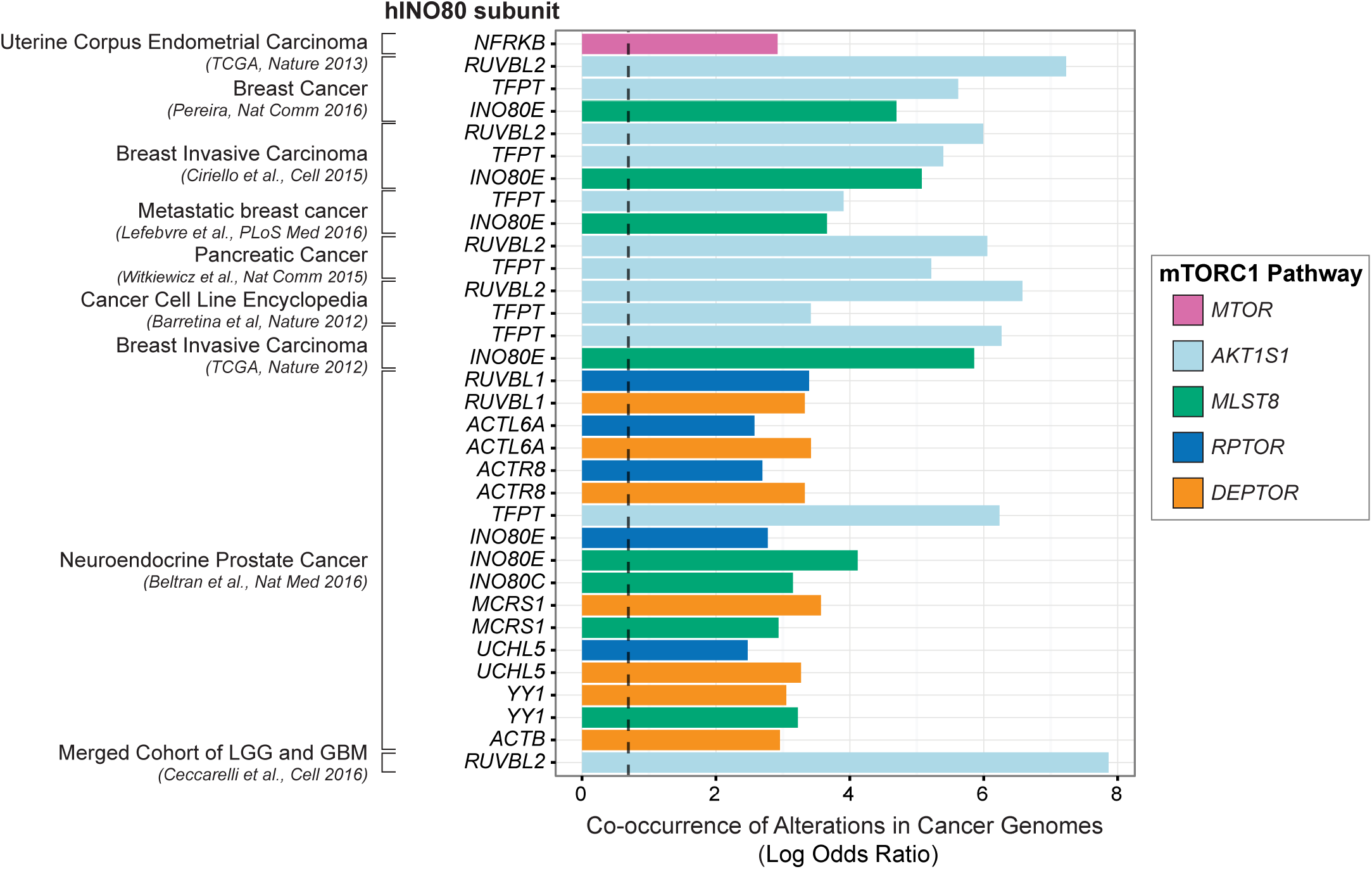
INO80 and mTORC1 alterations co-occur in cancers. (**D**) Co-occurrence of INO80 subunit and mTORC1 alteration in cancer using datasets from the cBioPortal (Cerami et al., 2012; Ciriello, Cerami, Sander, & Schultz, 2012; Gao et al., 2013). Datasets with high mutational load in the INO80/mTOR pathway gene sets (>20% altered samples) were used and small (< 50 samples) and provisional datasets were excluded. The natural log transformed odds ratio calculated by the mutual exclusivity tool in the portal is plotted for significant cooccurrences (Fisher’s Exact Test and false discovery rate of 0.001). Infinite calculated odds ratios are excluded. The dashed line marks tendency for co-occurrence (odds ratio of 2). Colors indicate mTORC1 subunits, human INO80 subunits are on the y-axis, co-occurrences are grouped by cancer study. The full table of significant co-occurrences is found in Supplementary File 8.

In summary, INO80, like many chromatin remodelers, has numerous roles in DNA-templated processes. Investigations of how these remodelers are controlled will likely reveal how chromatin modification is integrated with environmental responses. In this report, we have identified that the functions of the INO80 complex are modular, thus may be regulated in parts, rather than affecting the totality of INO80’s activity. Furthermore, we reveal that INO80 is involved in metabolic signaling, which likely contributes to adaptive gene expression responses in normal cells and may result in disease when disrupted.

## MATERIALS & METHODS

### Differential Genetic Interaction Screens

Genetic interaction screens (EMAPs) were performed as described (M Schuldiner, Collins, Weissman, & Krogan, 2006) except that the last selection step was performed by replica-plating cells on medium containing YPD (untreated), 10nM Rapamycin on SC, or YPD lacking glucose and containing 2% ethanol. Images for score calculations were taken 24 hours after pinning except for ethanol which was taken 48 hours afterwards. Static and differential genetic interaction scores were calculated using a MATLAB-based software toolbox as described (Bandyopadhyay et al., 2010; Collins et al., 2006) using standard significance thresholds for the static conditions (S ≥ 2.0 or S ≤ -2.5) and the differential conditions (S ≥ 3.0 or S ≤ -3.0).

### Yeast Strains

Yeast strains are listed in Supplementary File 9. Strain construction was in S288C background using standard techniques. All FLAG epitopes were chromosomally integrated to ensure endogenous expression of protein. Gene knockouts were integrated at the chromosomal locus.

The EMAP query strains are haploid Mat*α* yeast, as in Schuldiner et al., 2006, containing NAT marked mutations with the following background genotype: *his3Δ1 leu2Δ0 LYS2+ met15Δ0 ura3Δ0 can1Δ::MATa STE2Pr-HIS3 lyp1Δ::MATα STE3Pr-LEU2*. The EMAP test strains are haploid Mat*a* yeast, as in Ryan et al., 2012, containing KAN^R^ marked mutations with the following background genotype: *his3Δ1 leu2Δ0 met15Δ0 ura3Δ0. D*ecreased *a*bundance by *m*RNA *p*erturbation (DAmP) alleles are as previously described (Schuldiner et al., 2005).

### INO80 Subunit Domain Mutants

The following domain mutants of Ino80, Arp5, and Ies6 were used in this study. For the Ino80 ATPase subunit that scaffolds the complex, we deleted 3 domains: amino acids 2-200 (N-terminus, Nterm), which is required for association of the Nhp10 module (Ies1, Ies3, Nhp10, Ies5); the helicase-SANT-associated (HSA) domain (Szerlong et al., 2008) required for association of the Arp8 module (Arp8, Arp4, Act1, Ies4); and the insertion domain that splits Ino80’s two RecA ATPase lobes and is required for association of the Arp5 and Rvb1/2 modules (Arp5, Ies6, Ies2, Rvb1, Rvb2) (Yao et al., 2015). Two previously described domain mutants of the Arp5 subunit that are conserved across species but unique to Arp5 and help couple ATPase activity to productive nucleosome sliding (Yao et al., 2015) were used (D2 and D3).

For the Ies6 subunit that is a component of the Arp5 module (Yao et al., 2015), domain deletions across *IES6* based on conservation, hydrophobicity, intrinsic disorder, and protein interactions were created. We individually deleted two regions of the YL1-C domain, which is needed for the Arp6-Ies6 subcomplex to associate with INO80 (Yao et al., 2015). The C-terminal deletion (D5) strain was viable but EMAP results from this query did not pass quality control analysis and were subsequently excluded, while the N-terminal deletion (D4) query produced consistent EMAP results.

Domain mutants contain a C-terminal selectable marker after 500bp of endogenous 3’ sequence, except for the Swc2-YL1CΔ (AA 708-737Δ) mutant which has 449bp of 3’ sequence, and the Ino80-Nterm domain mutant, which contains a selectable marker 700bp upstream of the ORF, followed by endogenous 5’ sequence.

### Western Blotting

Protein from whole cell extracts were precipitated with 10% trichloroacetic acid. Proteins were detected by Western blot using anti-FLAG M2 (Sigma; catalog no. F1804), anti-hexokinase (Novus; catalog no. NB120-20547), anti-H3 C-terminal (Active Motif; catalog no. 39163), or anti-phospho-S6 ribosomal protein (Cell Signaling Technology; catalog no. 2211) antibodies.

### FLAG Affinity Purifications

Protein complexes were purified using FLAG affinity-agarose beads (Sigma; catalog no. A2220) as previously described (Yao et al., 2015), and washed with HEGN buffer containing 0.5M KCl.

### Bioinformatic Analysis

Bioinformatic analysis was conducted using R. Rankit normalization was performed using the formula (r – 0.5)/n (Solomon, 2008). Pearson correlations were performed using the cor() function, principal component analyses were performed using the prcomp() function. Genetic modules were determined using hierarchical clustering along with the kmeans() clustering algorithm. The number of centers was informed with a combination of a within sum of squares plot, average silhouette approach, and a gap statistic plot using the ‘factoextra’ R package, as well as a rational approach incorporating published structural data of the complex.

DAVID analysis was performed using version 6.7 with default parameters and medium stringency. Gene ontology (GO) enrichments were determined from GO annotations retrieved using the org.Sc.sgd.db R package (Bioconductor) after applying a multiple hypothesis corrected hypergeometric test using genes in the EMAP test library as background with a custom script.

Genome wide ChIP-seq correlations were performed using the Genome Track Analyzer (Kravatsky et al., 2015) on uniformly processed tracks using segment midpoints and considering both strands. The H3K56ac and Arp5 correlation reported in the text (r = 0.53) was calculated from uniformly processed data using the multiBigwigSummary and plotCorrelation tools in the deepTools2 suite (Ramírez et al., 2016) using 10 bp bins and Pearson correlation. Arp5 and H3K18ac occupancy profiles were generated from averaged ChIP traces ±1 kb around the +1 nucleosomes of all ORFs, smoothed by fitting a spline function selected by ordinary crossvalidation in R using smooth.spline(), then scaled and centered using the scale() function in R.

Network density was calculated as number of significant interactions observed divided by the total number of query-test gene pairs. Significance for network densities was assessed using a Monte Carlo randomization test. Randomization tests were performed with 100,000 permutations. Significance is notated as follows: * *p* < .05, ** *p* < .01, *** *p* < .001, n.s. is not significant.

### Petite Frequency Assay

Petite frequency was measured as previously described using a tetrazolium overlay (Hess et al., 2009).

### RNA-Sequencing

RNA was prepared from samples (approximately 1.5 ODs) using the MasterPure^™^ Yeast RNA Purification Kit (Epicentre, MPY03100). The sequencing libraries were prepared from 0.8 μg of RNA/sample using the Illumina TruSeq Stranded mRNA kit (Illumina, 15031047). The quality of the pooled library was checked using the Agilent Bioanalyser 2100 HS DNA assay. The library was sequenced on an Illumina HiSeq 2000 platform. Minimum of 10 million reads per sample were aligned using Bowtie 2 and analyzed using the DESeq2 package for R. Data deposition at NCBI is pending.

### Uniform ChIP-seq Processing

Reads were downloaded from GEO and uniformly processed. Briefly, reads were truncated to the smallest read length across datasets (36bp), mapped to the genome using STAR, and then signal coverage was generated and peaks were called using MACS2. Uniform processing of ChIP-seq data facilitates inter-study comparisons and can eliminate batch artifacts. Datasets used are listed in **Supplementary File 5**. +1 nucleosome positions were used as defined in (Jiang & Pugh, 2009). Datasets of insufficient quality after processing were excluded from subsequent analysis.

### ChIP-qPCR

ChIP was performed as previously described (Mizuguchi et al., 2004) with a few modifications. Cells were grown in YPD at 30 °C to OD_660_ of 0.7. Cells were lysed using Matrix D beads in a FastPrep homogenizer (MP Biomedicals) at maximum four times for 60 seconds, then sonicated to an average fragment size of 300 bp using a Bioruptor Plus (Diagenode) and clarified by centrifugation. Chromatin was immunoprecipitated using anti-H3K18ac (Millipore; catalog no. 07-354) pre-bound to Protein G Dynabeads (ThermoFisher; catalog no. 10004D) and washed 3 times in FA buffer with 150 mM NaCl then 2 times in FA buffer with 500 mM NaCl. DNA was eluted in TE with 1% SDS, cross-links were reversed by incubating overnight at 65 °C, treated with 0.2mg/ml RNAse A (VWR; catalog no. E866) for 2 hours at 37 °C, then extracted with phenol:chloroform:isoamylalchol and ethanol precipitated. DNA was resuspended in TE and analyzed by real-time quantitative PCR using iTaq Universal SYBR Green Supermix (BioRad; catalog no. 1725121). Ct values were determined using a CFX96 real-time detection system (BioRad).

## ACKNOWLEDGEMENTS

We wish to thank Stefan Bohn for technical advice with the EMAP and Nevan Krogan for providing the test library. We are grateful to members of the Morrison laboratory for helpful discussions. This work was supported by a Stanford Graduate Fellowship and NHGRI (5T32HG000044) to SLB, Coca-Cola Foundation Fellowship and Sr. Luis Alberto Vega Ricoy research support to PEGN, and NIH (R35GM119580) to AJM.

## AUTHOR CONTRIBUTIONS

SLB, Conceptualization, Software, Validation, Formal Analysis, Investigation, Resources, Writing – original draft, Writing – review & editing, Visualization; EKS, Investigation, Project administration; PEGN, Investigation, Formal Analysis; DAK, Investigation, Software, Formal Analysis; GJG, Investigation, Formal Analysis; KMW, Investigation; WY, Investigation; TLE, Investigation; APP, Investigation; EP, Investigation; LRL, Investigation; AJM, Conceptualization, Investigation, Writing – original draft, Writing – review & editing, Supervision, Project administration, Funding acquisition.

## COMPETING INTERESTS

The authors declare no competing interests.

---

**Figure 1 – figure supplement 1.**
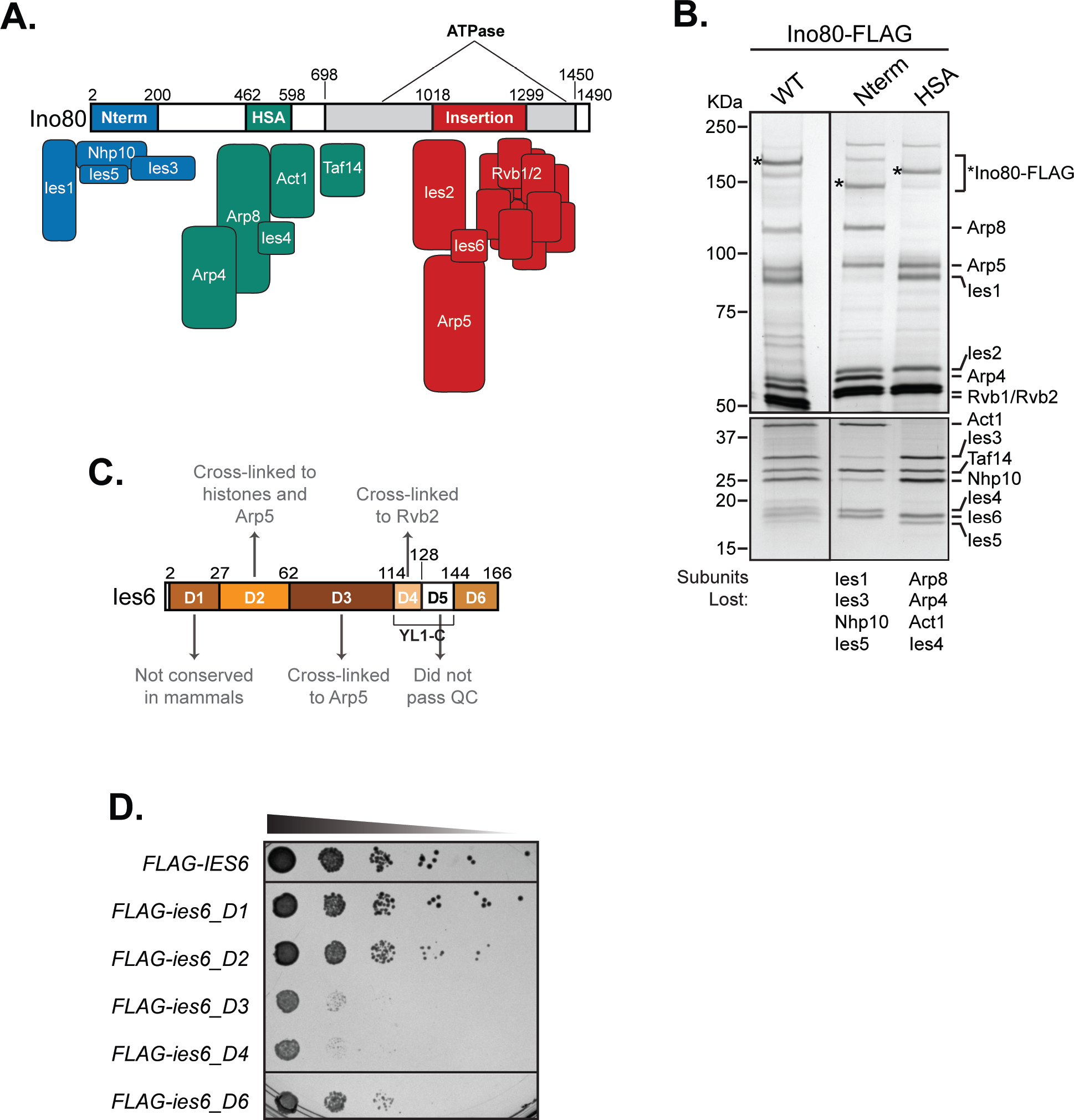
Ies6 and Ino80 domain mutants created for this study. (**A**) Schematic of the Ino80 ATPase protein domains with subunit binding modules illustrated from previous structural and biochemical studies (Tosi et al., 2013; Yao et al., 2015). Ino80 insertion is as described by (Ebbert, Birkmann, & Schüller, 1999), HSA domain is as identified in (Szerlong et al., 2008), N-terminus (Nterm) is amino acids 2-200. (**B**) Ino80-FLAG purifications from wild-type (WT), N-terminal deletion (Nterm), and HSA deletion strains were electrophoresed on 6% (top) and 15% (bottom) SDS-PAGE gels and identified by asterisk. Proteins were visualized via silver staining. Subunits of the INO80 complex are labeled on the right, molecular mass (KDa) is labeled on the left. Subunits lost from the INO80 complex are identified at the bottom. (**C**) Schematic of Ies6 gene domains, the YL1-C domain is split into domain 4 (D4) and domain 5 (D5). D5 was omitted from additional assays because EMAP results did not pass quality control (QC). (**D**) Fitness assay of indicated FLAG-tagged domain mutants described in (C). 1:10 serial dilution of strains were grown for 3 days at 30 °C on YPD.

**Figure 1 – figure supplement 2.**
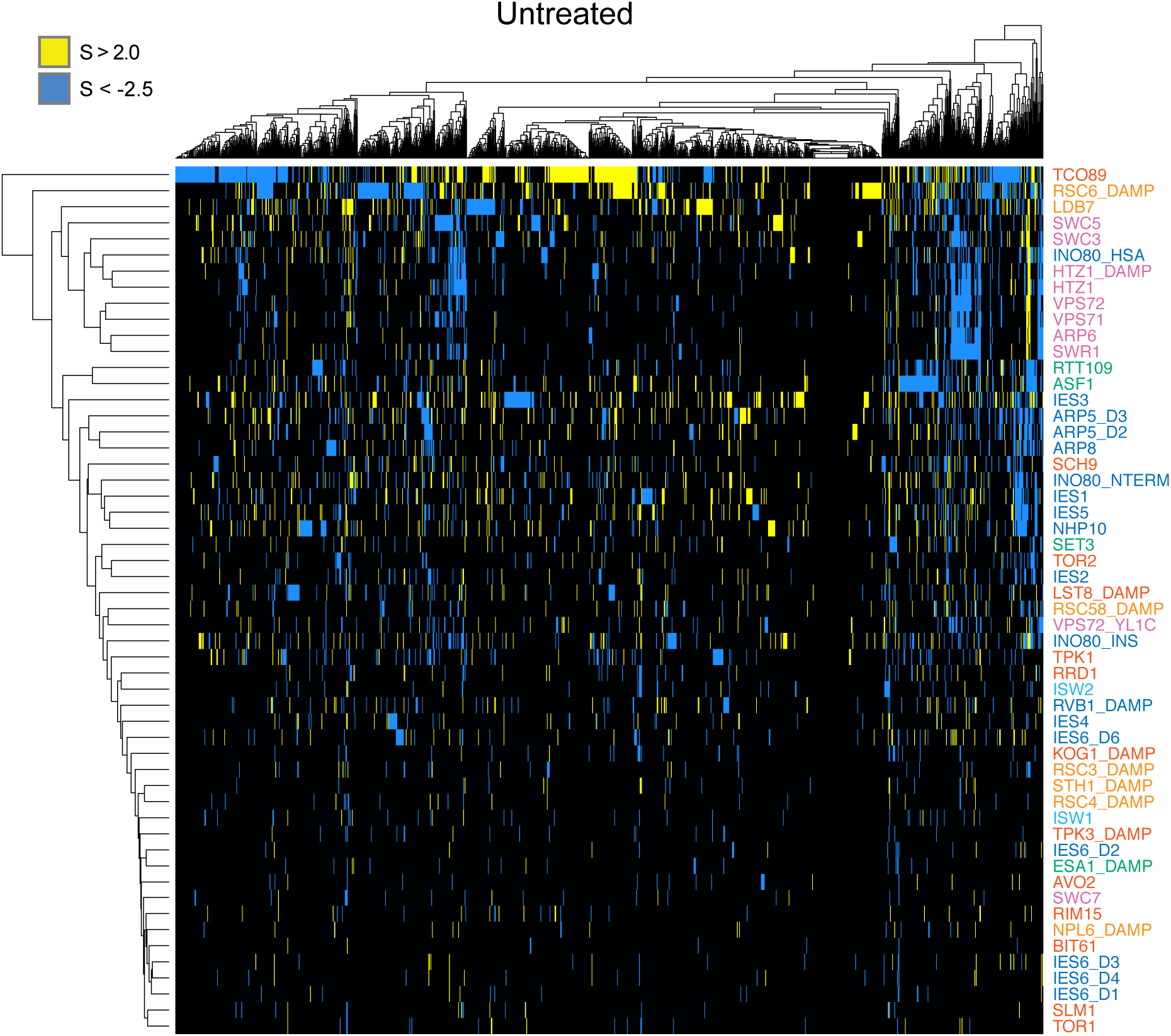
Hierarchical clustering of genetic interaction scores in untreated condition. Clustergram of all significant interactions in the untreated static EMAP between the 54 query strains and all test strains with at least one significant interaction. Test strains are along the x-axis. Text colors correspond to the query category annotated in Figure 1B. Supplementary File 1 lists all EMAP scores.

**Figure 1 – figure supplement 3.**
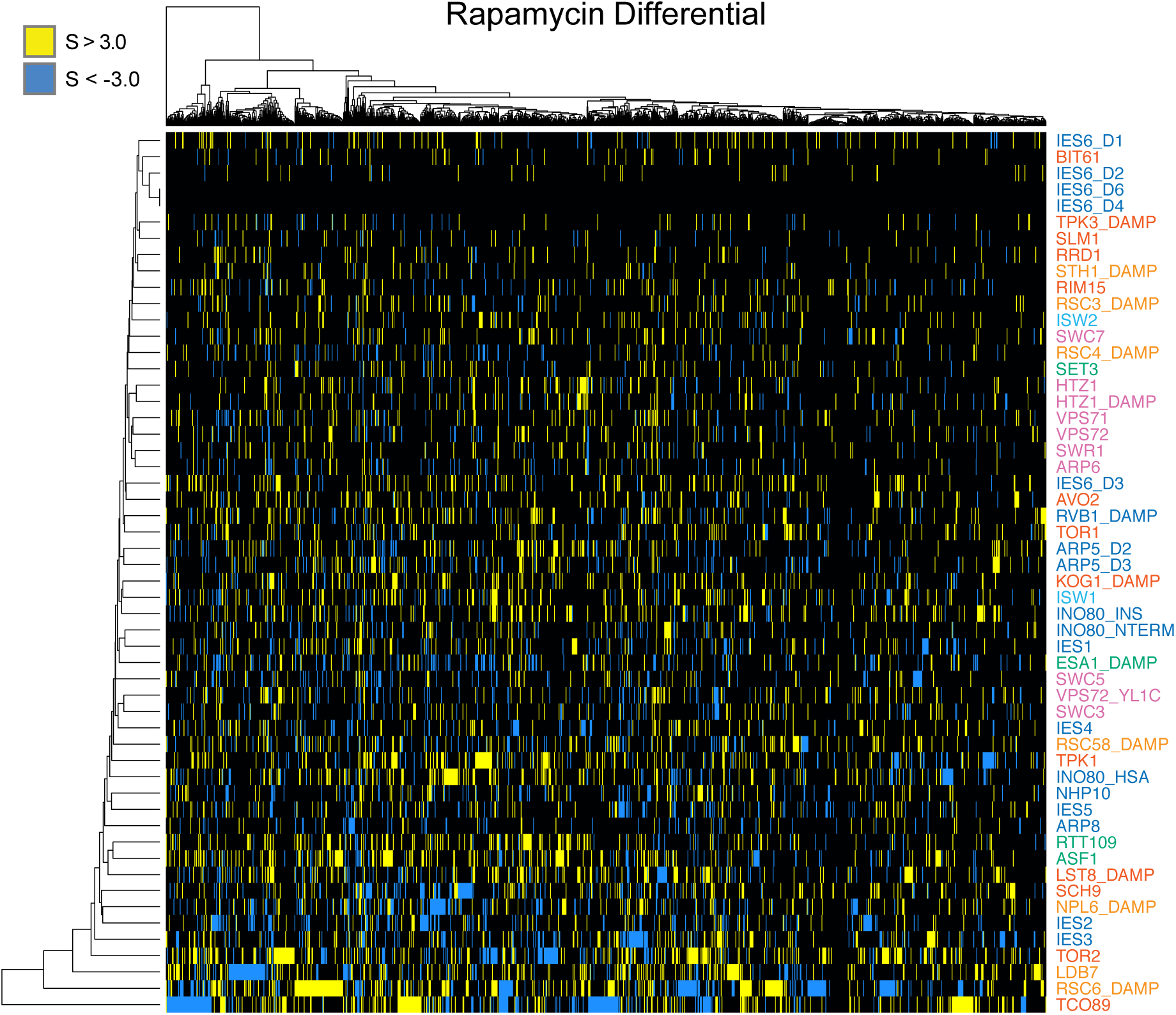
Hierarchical clustering of genetic interaction scores in the rapamycin differential condition. Clustergram of all significant interactions in the rapamycin differential EMAP between the 54 query strains and all test strains with at least one significant interaction. Test strains are along the x-axis. Text colors correspond to the query category annotated in Figure 1B. Supplementary File 1 lists all EMAP scores.

**Figure 1 – figure supplement 4.**
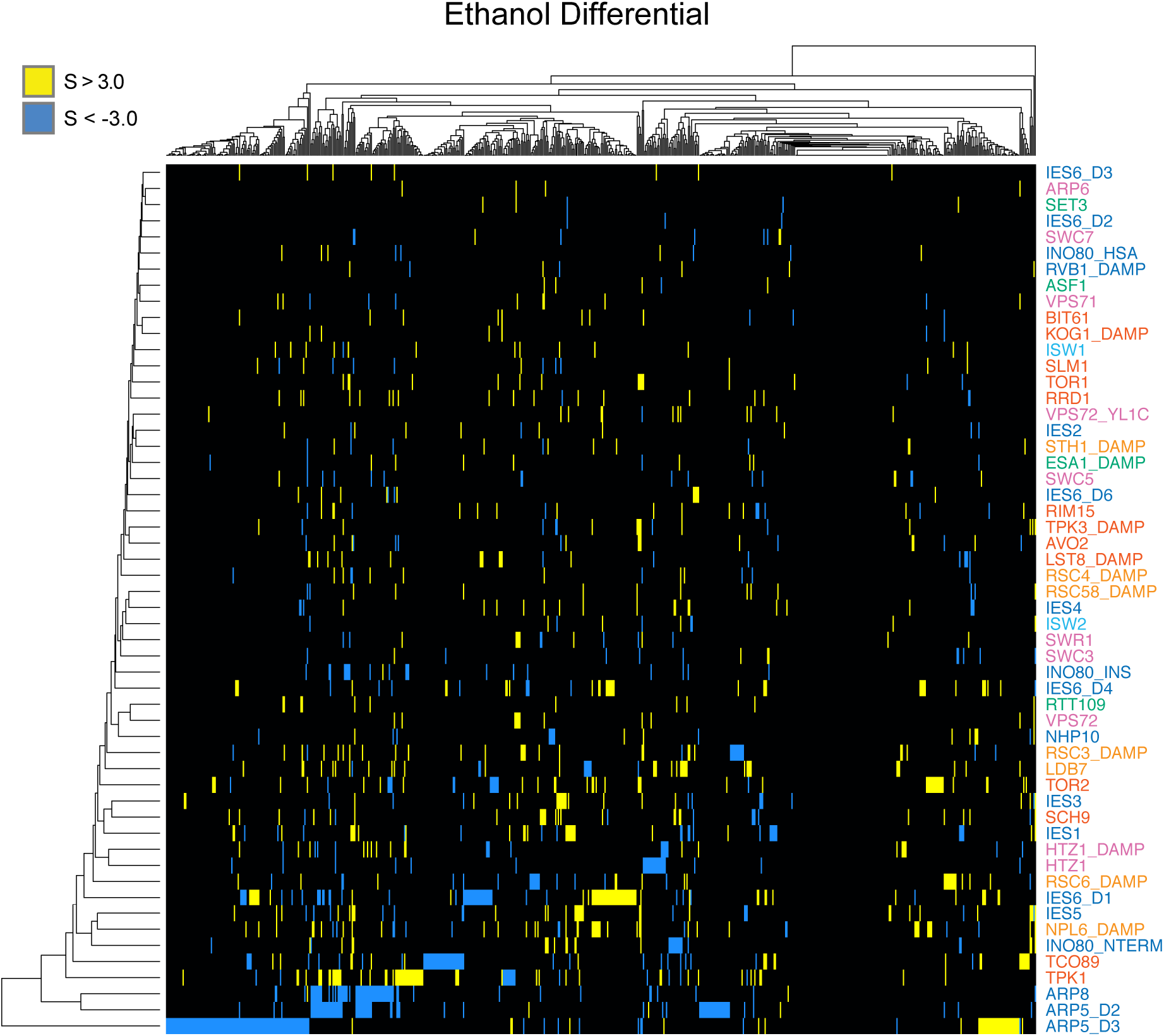
Hierarchical clustering of genetic interaction scores in the ethanol differential condition. Clustergram of all significant interactions in the ethanol differential EMAP between the 54 query strains and all test strains with at least one significant interaction. Test strains are along the x-axis. Text colors correspond to the query category annotated in Figure 1B. Supplementary File 1 lists all EMAP scores.

**Figure 1 – figure supplement 5.**
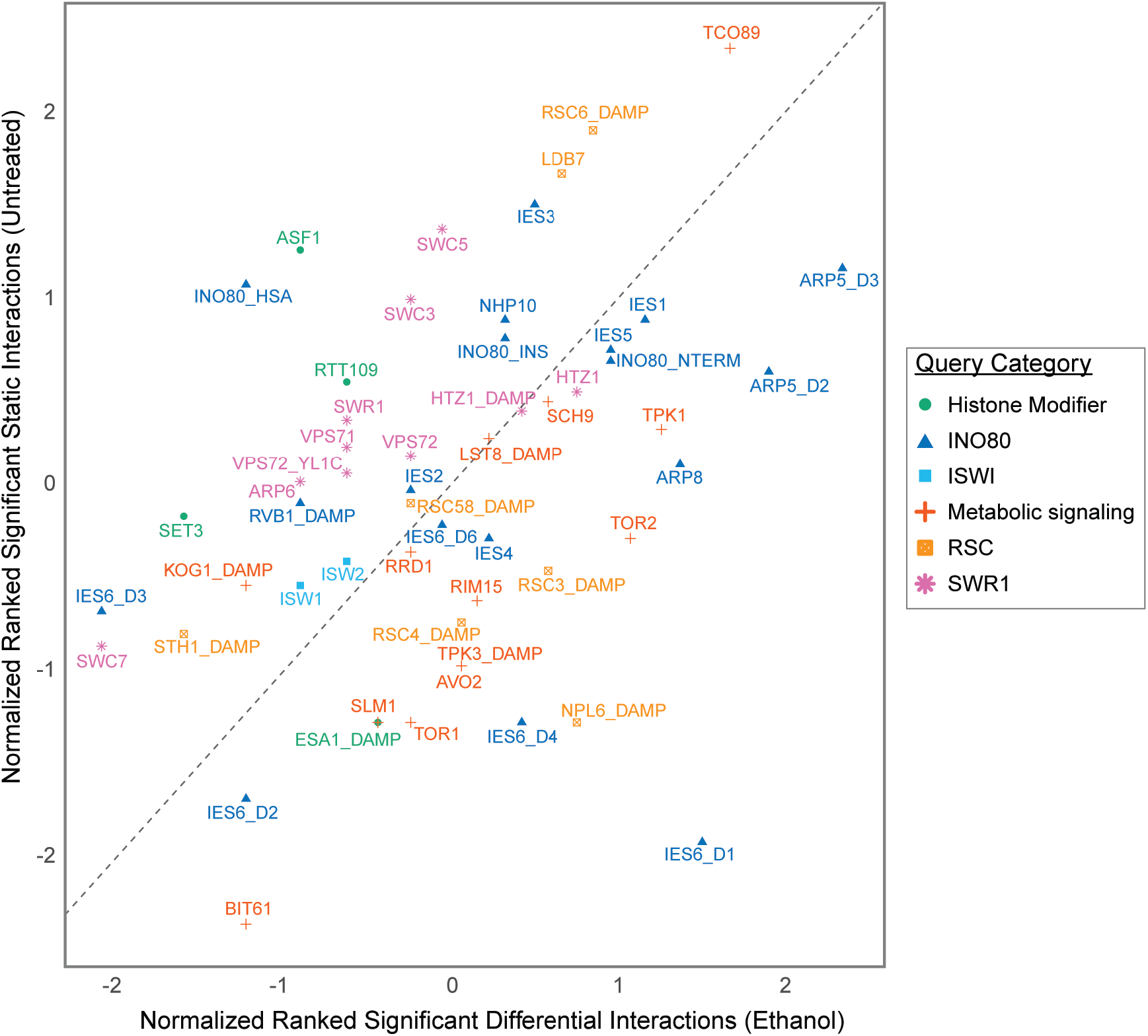
Significant interactions in the ethanol differential network. Plot of normalized significant interactions by query gene in the untreated condition and the ethanol differential condition, as in Figure 1F.

**Figure 2 – figure supplement 1.**
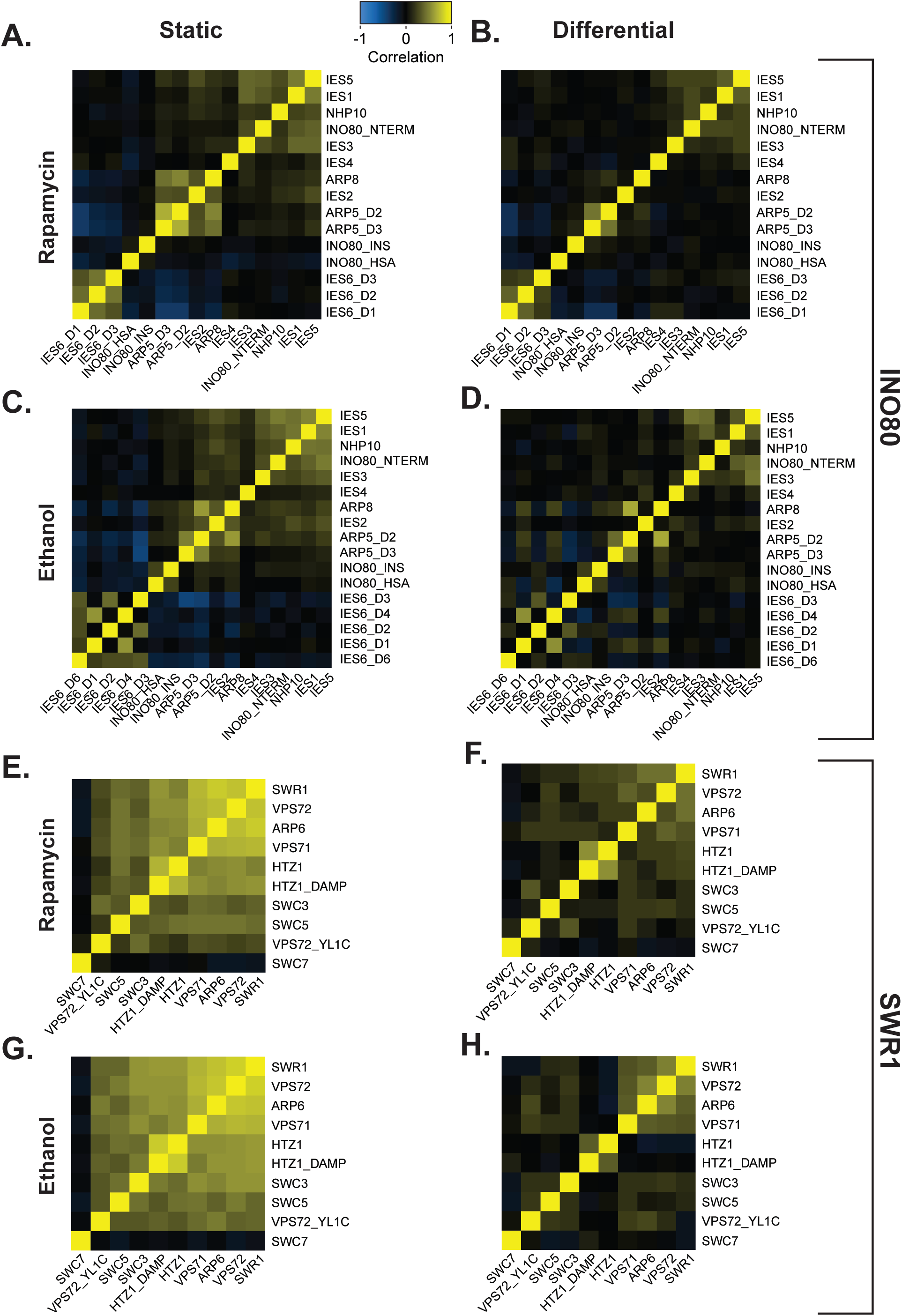
Genetic organization of INO80 and SWR1 in rapamycin and ethanol. Heatmap illustrating pairwise Pearson correlations between INO80 (A-D) and SWR1 (E-H) complex subunit query strains across the test library, as in Figure 2A. Rapamycin static correlations (A and E) and differential correlations (B and F) are shown. Ethanol static correlations (C and G) and differential correlations (D and H) are shown. Strains are ordered as shown in Figure 2A and determined by untreated hierarchical clustering.

**Figure 2 – figure supplement 2.**
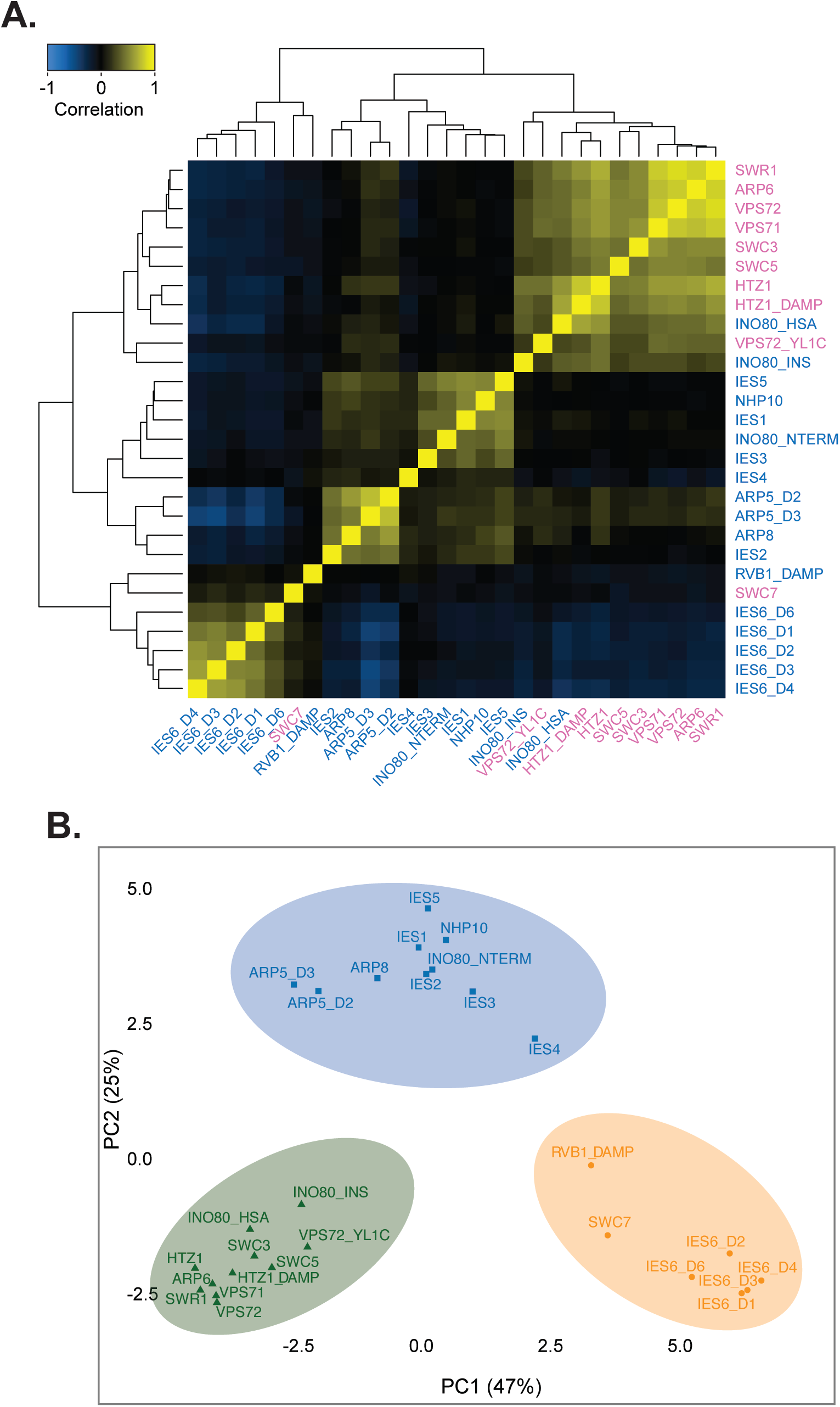
Genetic organization of the INO80 subfamily of remodeling complexes. (**A**) Heatmap of Pearson correlation of INO80 and SWR1 complex subunit query strains in the untreated static condition, as in Figure 2A; colors delineate complexes as in Figure 1B. Mutants are knockout or domain deletions where indicated: *INO80* N-terminal (NTERM), insertion (INS), and HSA deletions; *ARP5* domain 2 and 3 (D2 and D3) deletions; and *IES6* domain 1, 2, 3, 4, and 6 (D1, D2, D3, D4, D6) deletions. *D*ecreased *a*bundance by *m*RNA *p*erturbation (DAmP) alleles are as described in Schuldiner et al., 2005. Boxes outline subunit clusters identified by hierarchical clustering. (**B**) Principal component analysis (PCA) of INO80 and SWR1 complex subunit query strain Pearson correlations, as in Figure 2B. Colors indicate clusters identified by k-means clustering (k=3).

**Figure 5 – figure supplement 1.**
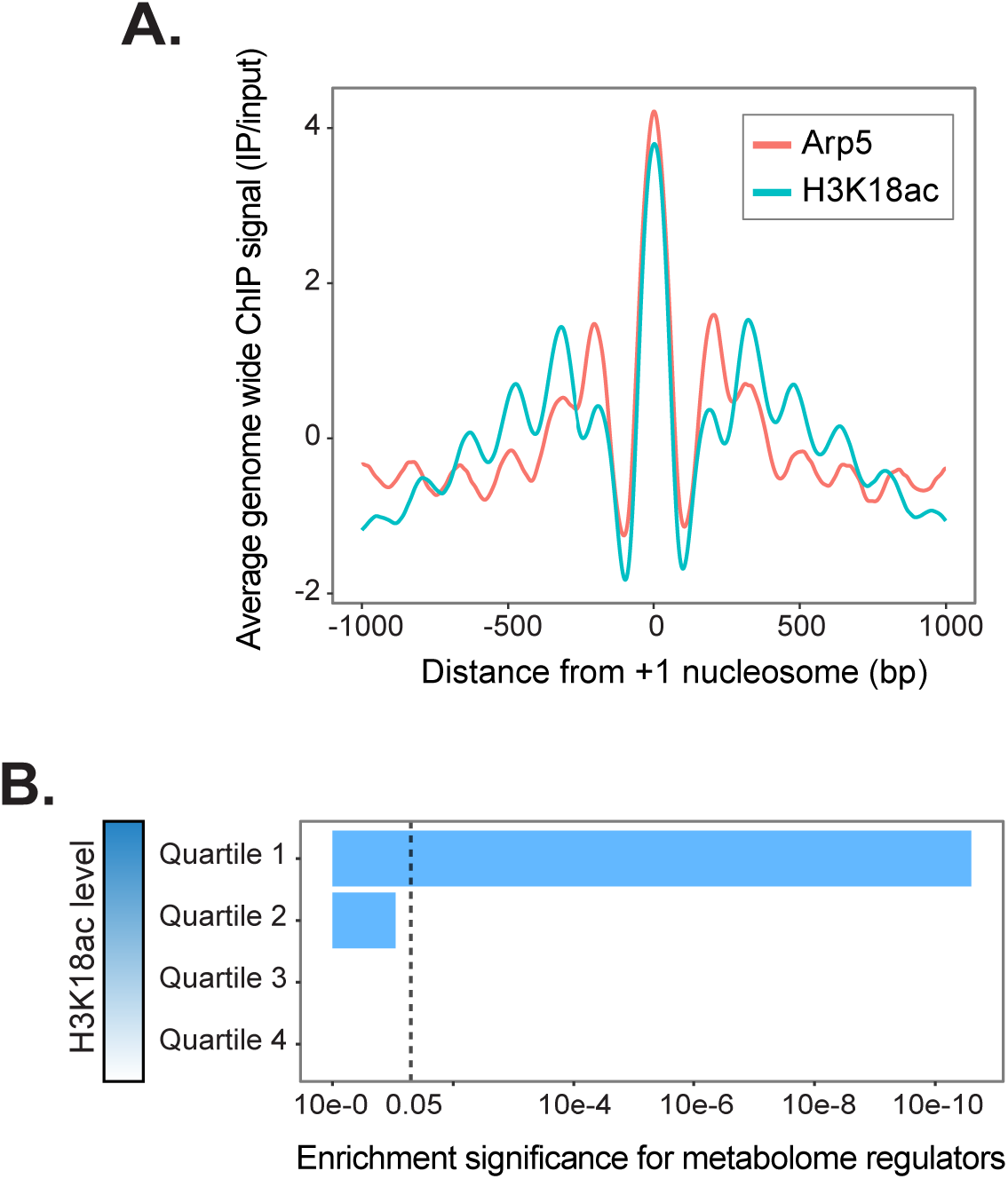
H3K18ac genome occupancy profile. (**A**) Genome-wide average uniformly processed (see *Materials and Methods*) ChIP-seq levels ±1000 bp from +1 nucleosomes (Jiang & Pugh, 2009) of Arp5 (Xue et al., 2015) and H3K18ac (Weiner et al., 2015). (**B**) Genes with high H3K18ac levels at +1 nucleosomes are significantly enriched for regulators of the metabolome; significance was determined using a hypergeometric test.

**Figure 5 – figure supplement 2.**
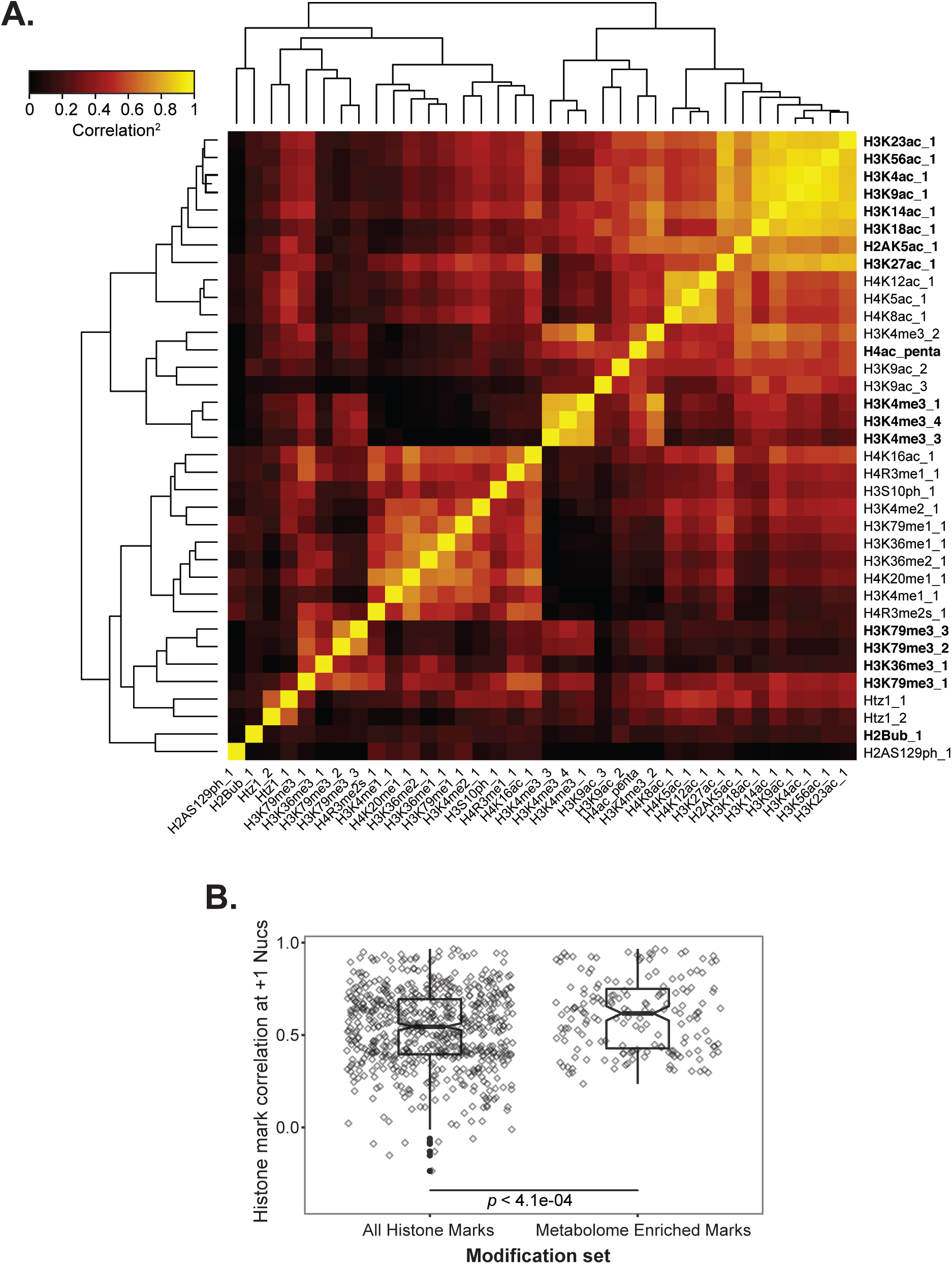
Histone modifications correlate with one another at +1 nucleosomes. (**A**) Heatmap of pairwise squared Pearson correlations at +1 nucleosomes using uniformly processed published ChIP-seq data (see *Materials and Methods*). Modifications that have significantly high levels at the +1 nucleosomes of metabolome regulators and are enriched for metabolome regulators in their top quartile of +1 nucleosome levels are bolded. (**B**) Box and jittered scatter plots of correlations between all histone marks shown and the metabolome enriched marks bolded in (A). Significance is determined using a Wilcoxon rank sum test (*p* < 4.1e-4) and by Monte Carlo randomization test (*p* = 0.0419).

**Figure 5 – figure supplement 3.**
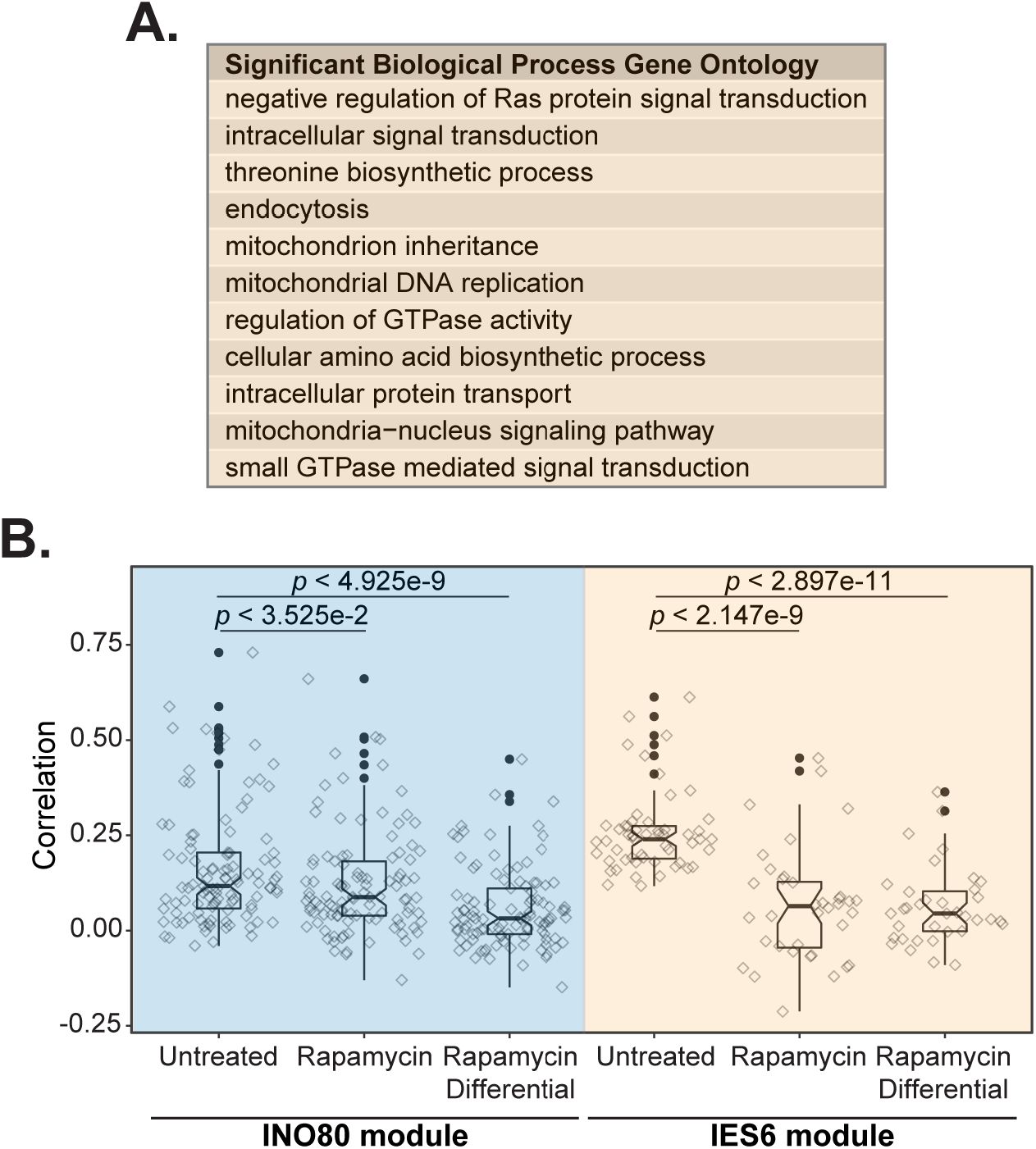
IES6 clusters in a rapamycin-sensitive metabolic module. (**A**) Table showing select gene ontology (GO) terms enriched in test strains that significantly interact with the IES6 cluster query genes (FDR-adjusted hypergeometric test, *p* < .05). The complete list of significant GO terms is found in Supplementary File 6. (**B**) Box and jittered scatter plots of correlations between query genes in the INO80 and IES6 expanded modules, shown in Figure 4A, in the untreated static, rapamycin static and differential conditions. Significance is determined using a Wilcoxon rank sum test.

**Figure 6 – figure supplement 1.**
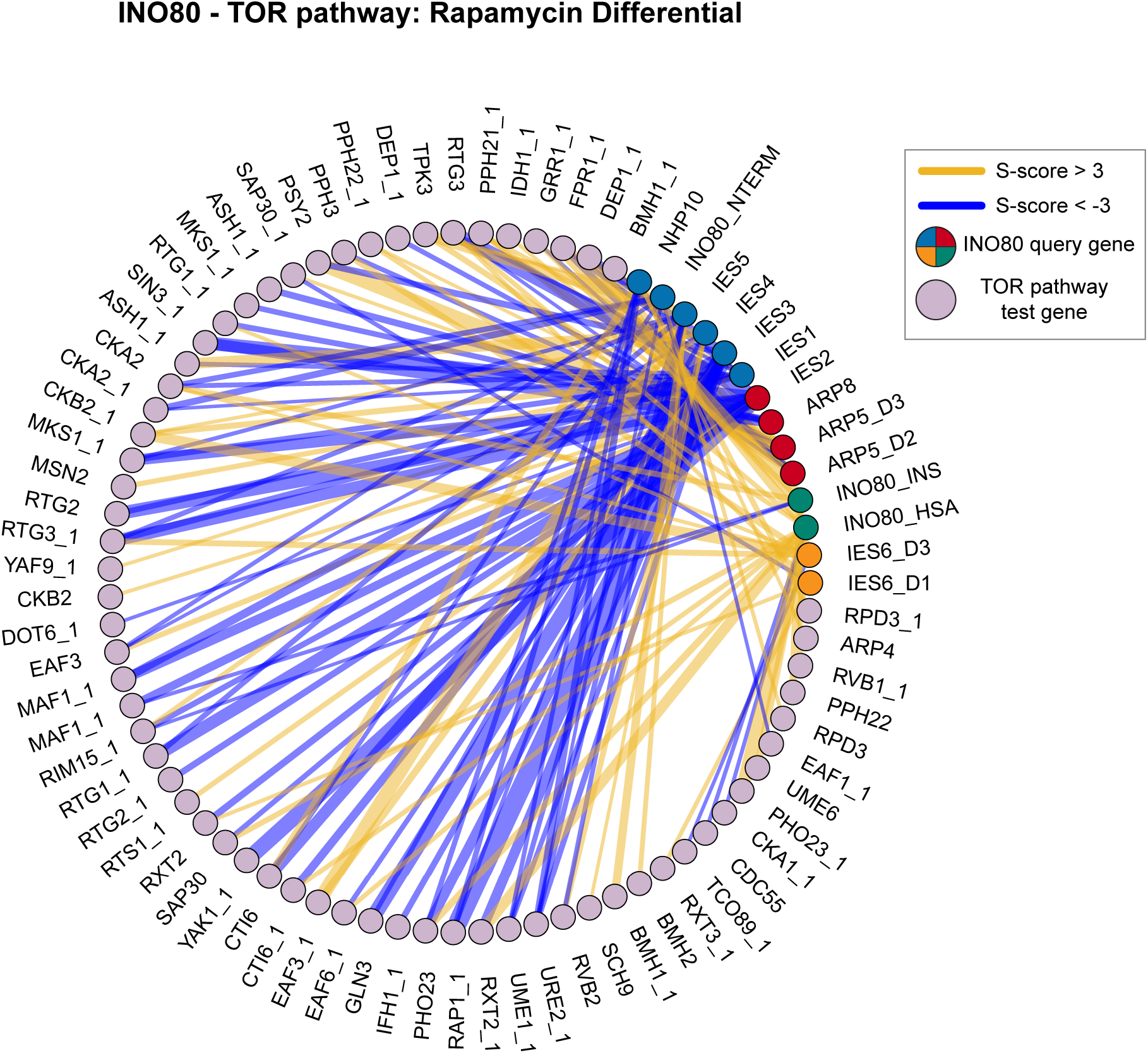
INO80 and the TOR pathway have a highly connected genetic interaction network in the rapamycin EMAP. Genetic interaction network between INO80 subunit query strains and significantly interacting TOR pathway test strains in the rapamycin differential condition. Line width indicates strength of S-score, INO80 queries are colored according to modules identified in Figure 2. Network density is significantly high, *p*-value = 1.6e-4 by Monte Carlo randomization test.

## ADDITIONAL FILES

**Supplementary File 1**. Genetic interaction data generated by static and differential EMAPs.

**Supplementary File 2**. Significant query interactions by treatment, related to Figure 1.

**Supplementary File 3**. DAVID functional annotation clusters by module, related to Figure 3.

**Supplementary File 4**. Gene ontology enrichments by module, related to Figure 3.

**Supplementary File 5**. ChIP-seq datasets uniformly processed for analysis in this study, related to Figures 4 and 5.

**Supplementary File 6**. Gene ontology enrichments of the expanded IES6 genetic module, related to Figure 6.

**Supplementary File 7**. Lists of genes in pathways utilized in this study, related to Figures 6, 7, and 8.

**Supplementary File 8**. Mutual exclusivity data generated by cBioPortal for INO80 and mTORC1 subunits, related to Figure 9.

**Supplementary File 9**. (A) Yeast strains used in this study. (B) EMAP query strains used in this study. (C) EMAP test strains used in this study.

